# High-throughput sequencing-based neutralization assay reveals how repeated vaccinations impact titers to recent human H1N1 influenza strains

**DOI:** 10.1101/2024.03.08.584176

**Authors:** Andrea N. Loes, Rosario Araceli L. Tarabi, John Huddleston, Lisa Touyon, Sook San Wong, Samuel M. S. Cheng, Nancy H.L. Leung, William W. Hannon, Trevor Bedford, Sarah Cobey, Benjamin J. Cowling, Jesse D. Bloom

**Affiliations:** Howard Hughes Medical Institute, Seattle, WA; Division of Basic Sciences, Computational Biology Program, and Vaccine and Infectious Disease Division, Fred Hutchinson Cancer Center, Seattle, WA; HKU-Pasteur Research Pole, School of Public Health, The University of Hong Kong, Hong Kong, SAR, China; World Health Organization Collaborating Centre for Infectious Disease Epidemiology and Control, School of Public Health, The University of Hong Kong, Hong Kong, SAR, China; Molecular and Cellular Biology Graduate Program, University of Washington, Seattle, WA 98109, USA; Department of Ecology and Evolution, University of Chicago, Chicago, IL

## Abstract

The high genetic diversity of influenza viruses means that traditional serological assays have too low throughput to measure serum antibody neutralization titers against all relevant strains. To overcome this challenge, we have developed a sequencing-based neutralization assay that simultaneously measures titers against many viral strains using small serum volumes via a workflow similar to traditional neutralization assays. The key innovation is to incorporate unique nucleotide barcodes into the hemagglutinin (HA) genomic segment, and then pool viruses with numerous different barcoded HA variants and quantify infectivity of all of them simultaneously using next-generation sequencing. With this approach, a single researcher performed the equivalent of 2,880 traditional neutralization assays (80 serum samples against 36 viral strains) in approximately one month. We applied the sequencing-based assay to quantify the impact of influenza vaccination on neutralization titers against recent human H1N1 strains for individuals who had or had not also received a vaccine in the previous year. We found that the viral strain specificities of the neutralizing antibodies elicited by vaccination vary among individuals, and that vaccination induced a smaller increase in titers for individuals who had also received a vaccine the previous year—although the titers six months after vaccination were similar in individuals with and without the previous-year vaccination. We also identified a subset of individuals with low titers to a subclade of recent H1N1 even after vaccination. This study demonstrates the utility of high-throughput sequencing-based neutralization assays that enable titers to be simultaneously measured against many different viral strains. We provide a detailed experimental protocol (DOI: https://dx.doi.org/10.17504/protocols.io.kqdg3xdmpg25/v1) and a computational pipeline (https://github.com/jbloomlab/seqneut-pipeline) for the sequencing-based neutralization assays to facilitate the use of this method by others.

## Introduction

Human influenza virus evolves rapidly to escape immunity. Newly evolved strains often possess mutations in the viral hemagglutinin (HA) surface protein that erode neutralization by human polyclonal antibodies (1). In general, protection against infection is correlated with serum antibody neutralization and hemagglutination-inhibition (HAI) titers (2–4), and viral strains with reduced susceptibility to neutralization by human serum antibodies are more likely to spread widely (5).

Unfortunately, traditional methods for measuring neutralization or HAI titers are typically unable to provide information on the full swath of relevant viral diversity. The reason is that traditional neutralization or HAI assays test a single serum sample against a single viral strain in each measurement, meaning that vast numbers of assays (as well as large serum volumes and virus stocks) are needed to characterize titers against many different viral strains. This is a serious limitation, as the strain (or strains) chosen to test in the neutralization or HAI assays will often differ from the strain to which any given individual is exposed in any given year. For example, Petrie et al. found that HAI titer against the specific mutant that circulated that season was predictive of protection, but titer against the vaccine strain was not (6). Therefore, it would be far more informative if we could measure neutralization against the full influenza strain diversity to which an individual might be exposed, rather than just a few prototype reference strains.

Here we describe a high-throughput sequencing-based neutralization assay that simultaneously measures titers against many influenza virus strains. We applied this method to address an important question in influenza immunity (7–10): namely, how does repeated vaccination affect antibody titers? We found substantial variation among individuals in neutralization titers against recent human H1N1 strains. In addition, consistent with earlier studies (8,9,11,12), we found that individuals who received the vaccine in the previous season had a smaller increase in neutralization titers than individuals not vaccinated in the previous season. However, despite this smaller titer increase in repeat-vaccinees, the median absolute titers six months post vaccination were similar between singly and repeat-vaccinated individuals. Finally, we identified an antigenically distinct clade of human H1N1 influenza to which some individuals show low titers post vaccination. Overall, this work illustrates the power of high-throughput sequencing-based assays to map neutralization landscapes against a broad swath of influenza strains.

## Results

### Sequencing-based strategy for simultaneously measuring neutralizing titers against dozens of viruses

Our goal was to measure titers against many influenza viruses using the same workflow as a traditional neutralization assay. The fundamental idea behind our approach is to pool many viruses each encoding a different HA, and then read out neutralization of all these viruses simultaneously using next-generation sequencing (Fig. 1). Implementation of this approach required two key technical innovations. First, we incorporated a unique 16-nucleotide barcode sequence within the genomic segment encoding the HA of each strain (13,14), thereby allowing each HA to be identified by sequencing just this short barcode region (Fig. S1). Second, we developed a spike-in control RNA to enable conversion of sequencing counts of the viral barcodes to fraction viral infectivity retained upon serum treatment (Fig. 1).

**Figure 1.**
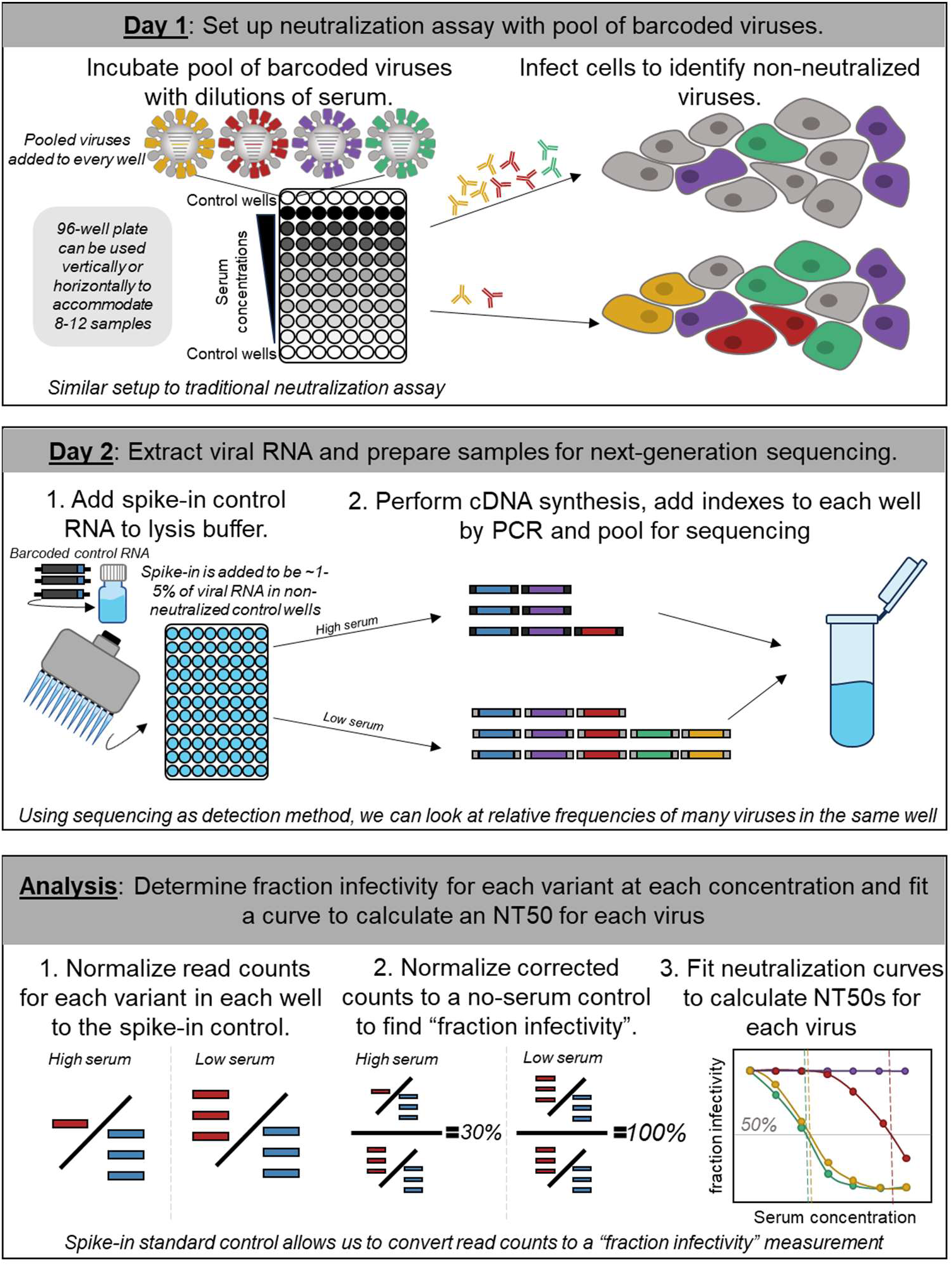
Approach for sequencing-based neutralization assay. A pool of barcoded viruses is incubated with serially diluted serum. Cells are added to the virus-serum mixes, and non-neutralized viruses infect cells where they produce viral RNA. At 16 hours post infection, lysis buffer supplemented with an RNA spike-in control is added to each well, and the RNA is extracted and processed for next-generation sequencing. To calculate the fraction infectivity of each virus at each serum concentration from the sequencing data, the sequencing counts for each viral barcode are first normalized by the counts of the spike-in control in each well, and then further normalized by the same ratio in no-serum control wells. The neutralization titers (NT50s) for each barcoded virus are calculated by fitting a Hill curve to the fraction infectivity measurements.

More specifically, we first created a library of influenza viruses encoding barcoded HAs from different natural strains (Fig. 1, S1). To perform the sequencing-based neutralization assays, we preincubated the library of barcoded viruses with serial dilutions of serum in a 96-well plate (Fig. 1). We then added cells to each well. The non-neutralized viruses infect cells and express high levels of their viral RNA, including the barcoded HA genomic segment. We extracted RNA from the cells and amplified the barcode region of the HA segment using PCR for next-generation sequencing. To quantify the absolute amount of barcoded viral RNA by sequencing, we added a constant amount of a spike-in control containing known barcodes (Fig. S1C), and normalized the viral barcode counts by the counts for the spike-in control in each well (Fig. 1). We quantified the fraction infectivity remaining for each barcoded virus at each serum concentration as the ratio of the normalized counts of that barcode to the normalized counts of that barcode in the no-serum control wells. We then fit neutralization curves to the fraction infectivity values, quantifying the neutralization titer 50% (NT50) as the serum concentration at which 50% of the viral infectivity is neutralized. With this approach, each row or column of the 96-well plate yielded neutralization titers against all the viral variants in the library. Because we designed the libraries to contain multiple distinct barcodes for each different HA, each row or column also yielded multiple replicate titer measurements for each virus. A more detailed description is provided in the methods and a step-by-step protocol is available on <u>protocols.io</u> (DOI: https://dx.doi.org/10.17504/protocols.io.kqdg3xdmpg25/v1). See https://github.com/jbloomlab/seqneut-pipeline for an open-source computational pipeline for analyzing the data.

### Library of barcoded viruses with HAs from recent human H1N1 strains

We aimed to use the sequencing-based assay to measure neutralization titers against the HAs of H1N1 influenza virus strains that circulated in humans between 2020-2022. Therefore, we designed a library comprised of the HAs of the five most recent human H1N1 vaccine strains and an additional 31 recent strains that were selected as representatives of subclades within a phylogeny constructed from sequences sampled over a 2-year timeframe, separated from their parent clade by at least one HA1 mutation, and detected at >1% globally (Fig. 2A). The HA ectodomains from the 31 recent strains and the two most recent vaccine strains included in the library vary by 52 amino-acid substitutions at 48 different sites (Fig. 2B), with most of the variable sites located in the globular head of HA in regions that are known to be antigenically important (15). There are additional sites that differ in the three older vaccine strains included in the library.

**Figure 2.**
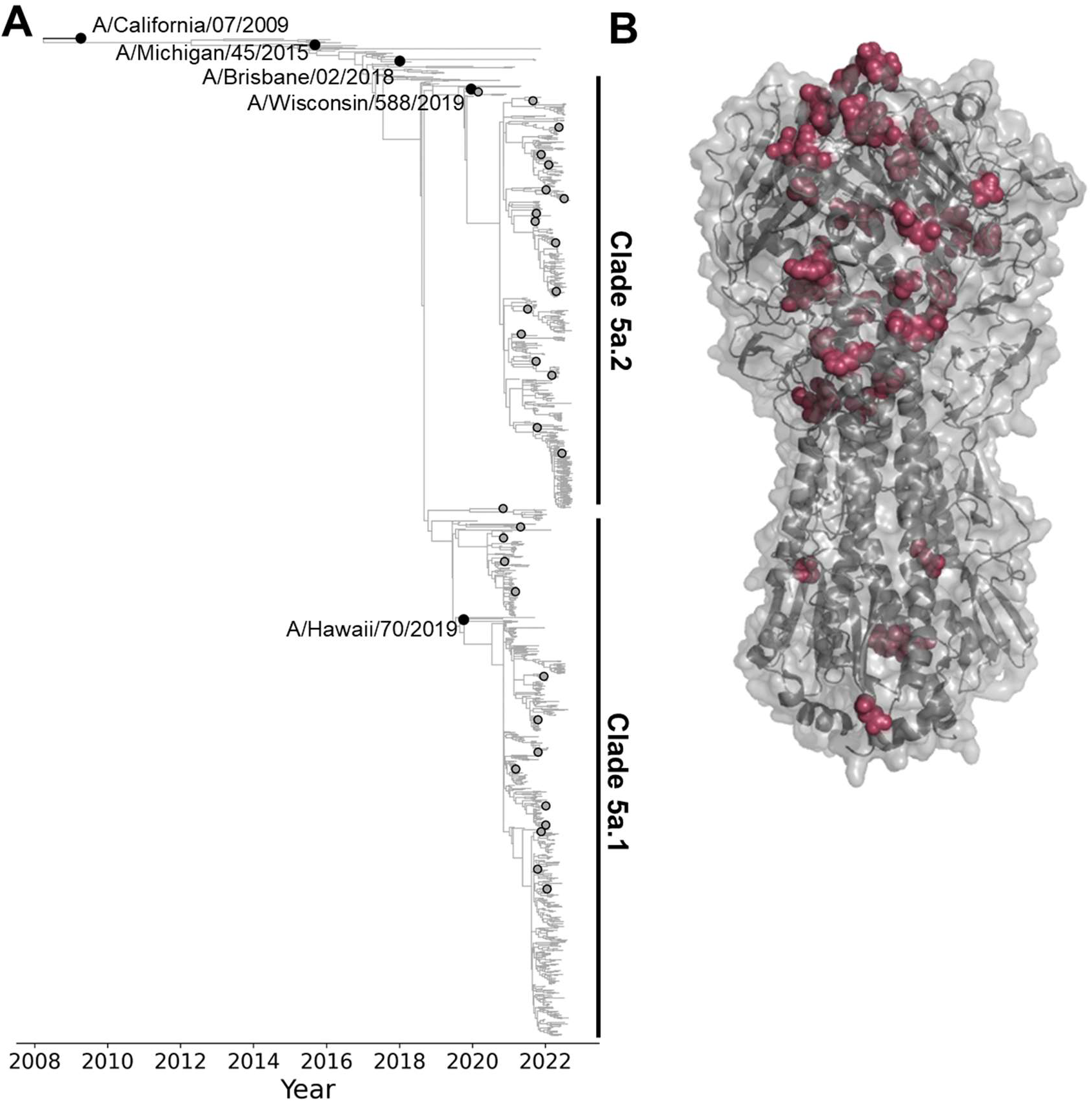
Human H1N1 influenza strains with HAs incorporated in the neutralization-assay library. A) Phylogeny of recent human H1N1 HA generated by nextstrain.org from October 2022 (56,57), with black circles indicating recent cell-based vaccine strains included in the library, and gray circles indicating strains selected for the library from recent emerging clades. The recent strains as well as the two most-recent vaccine strains (A/Hawaii/70/2019 and A/Wisconsin/588/2019) fall into the 5a.1 or 5a.2 clades as indicated on the tree. See https://nextstrain.org/flu/seasonal/h1n1pdm/ha/2y@2022-10-07?d=tree&p=fullhttps://nextstrain.org/flu/seasonal/h1n1pdm/ha/2y@2022-10-07?d=tree&p=full for an interactive version of the tree. B) Structure of HA indicating amino-acid sites that vary among the HA ectodomains of the 31 recent strains and the two most-recent vaccine strains included in the library (PDB: 6XGC) (70). Additional sites not indicated on the structure differ in the three older vaccine strains included in the library. The overall HA structure is shown in gray, and sites that vary among recent library strains are shown as maroon spheres.

To generate the barcoded library, we cloned the ectodomain of each HA into a unidirectional influenza reverse genetics plasmid (16) that included a 16-nucleotide barcode introduced in a way that does not disrupt viral genome packaging (Fig. S1) (13,14). We generated plasmids with two to four distinct barcodes for each different HA to provide replicates within the library.

Therefore, although our library contained 36 unique HAs, there were 110 different barcodes in the library. We pooled the plasmids encoding replicate barcodes for the HA for each viral strain and then used reverse-genetics to separately generate each of the 36 small pools of two to four barcoded viruses encoding each HA. The seven non-HA influenza segments in these viruses were derived from the lab-adapted A/WSN/1933 (H1N1) strain (17), a choice that was made both for biosafety reasons and to ensure that all viruses differed only in their HAs and were otherwise identical. Note, although human serum contains antibodies to a variety of influenza proteins (18), typically only antibodies to HA are strongly neutralizing in single-cycle infection assays like the ones used in our experiments (19,20). The independently generated stocks of barcoded viruses with each different HA were then pooled so that each HA was represented at approximately equal transcriptional titers. To avoid separately titering dozens of viruses by TCID50 or plaque assays, we used a sequencing-based titering method (Fig. S2). Specifically, we first infected cells with an equal-volume mixture of all virus stocks, then used next-generation sequencing of the viral barcodes to determine the relative transcriptional output of each virus stock, and finally re-pooled to equalize these transcriptional titers across HAs, such that each strain contributed roughly the same fraction of transcribed barcodes when cells were infected with the re-pooled library in the absence of serum antibodies (Fig. S2).

### Example results from sequencing-based neutralization assay, and validation with traditional neutralization assays

Example neutralization curves from the sequencing-based neutralization assay against the H1N1 vaccine strain from 2021-2022 (A/Wisconsin/588/2019) are shown in Fig. 3A for serum samples collected from an individual at 0- and 30-days post vaccination. Note that each row of the 96-well plate generates curves like those in Fig. 3A for each of the 36 different viruses (Fig. S3), with each curve generated in replicate because there are multiple barcodes associated with each viral HA (three barcodes in the case of the virus in Fig. 3A). The curves are conceptually identical to those generated by a traditional neutralization assay, with a fraction viral infectivity measurement made for each serial dilution of serum, and a sigmoidal Hill curve fit to the data to estimate the serum dilution at which half of the viral infectivity is retained.

**Figure 3.**
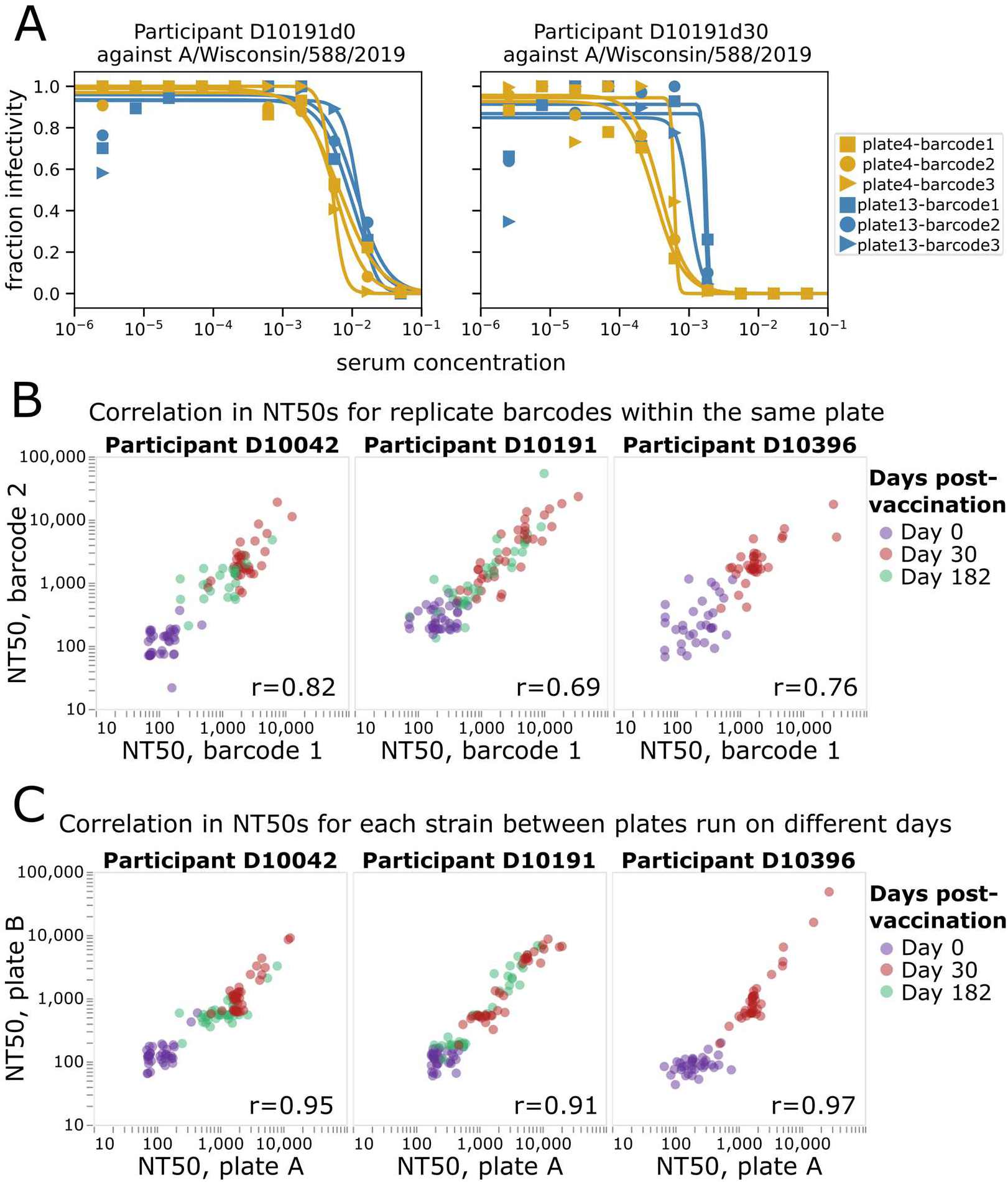
Sequencing-based neutralization assay yields reproducible measurements. A) Example sequencing-based neutralization assay curves for virus with the HA from A/Wisconsin/588/2019 (H1N1) against serum samples collected from study participant D10191 on day 0 or day 30 post vaccination. The library contains three different barcodes for this HA, so three curves (one for each barcode) are generated on each of two different plates (plate 4 and plate 13) that were run on different dates. See Fig. S3 for a display of all curves generated from a single plate. B) Correlation between NT50s measured for replicate barcodes corresponding to the same virus strain on the same plate. Each point represents a different viral strain, and each plot facet represents a different individual, with the colors indicating the days post vaccination the serum was collected from that individual. The Pearson correlation (r) is indicated on the plot. C) Correlation between NT50s measured for each viral strain on two different plates run on different days. The NT50s for each plate represent the median across the replicate barcodes for each virus on that plate.

The neutralization curves generated by the sequencing-based assay are highly reproducible. Reproducibility can be assessed at two levels: how similar are measurements for different barcodes associated with the same HA in each row (or column) of a single plate, and how similar are measurements made for the same HA across different plates run on different days. Visual inspection of Fig. 3A shows the consistency of curves both across barcodes within a row of a single plate and across different plates run on different days. A more systematic analysis of additional sera and viruses shows consistently high reproducibility of the NT50s measured both for replicate barcodes within a plate row (Fig. 3B) and the median NT50s measured for the same sera and viruses in different plates run on different days (Fig. 3C).

The neutralization curves and NT50s measured by the sequencing-based neutralization assay are very similar to those measured using a more traditional fluorescence-based one-virus versus one-serum single-cycle neutralization assay (21,22) (Fig. 4A, 4B). The sequencing-based neutralization assay NT50s also correlate well with HAI measurements for the 2020-2021 H1N1 vaccine strain, A/Hawaii/70/2019 (Fig. 4C). Note that the HAI titers are not expected to necessarily be identical to the neutralization titers, since HAI only assesses the ability of serum antibodies to prevent HA binding to sialic acid on red blood cells, whereas virus neutralization can also be caused by additional mechanisms in addition to blocking of receptor binding (23–25).

**Figure 4.**
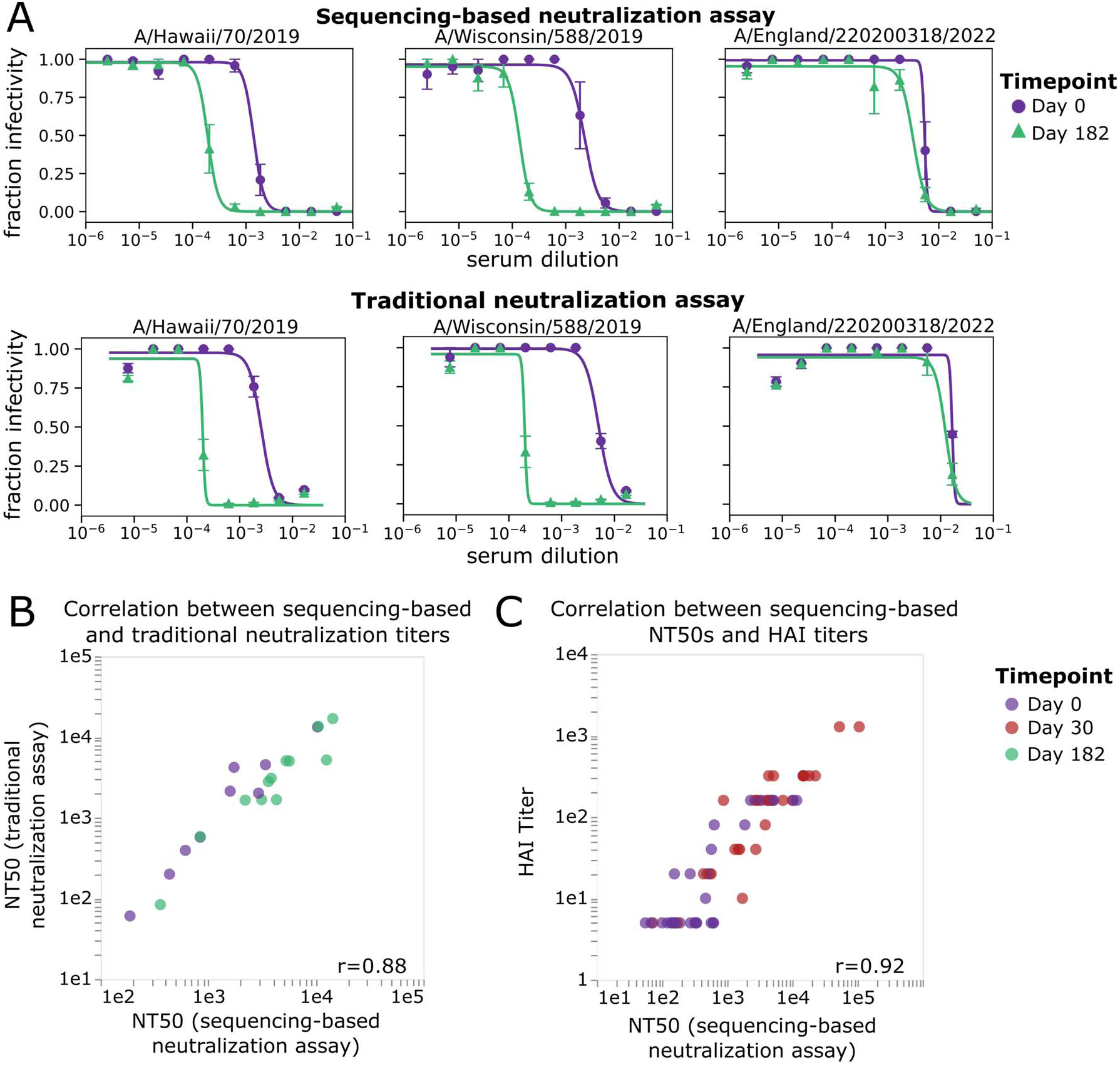
NT50s measured with the sequencing-based neutralization assay are highly correlated with those measured using a traditional neutralization assay. A) Example neutralization curves against sera collected from study participant D10378 collected on day 0 or day 182 post vaccination, measured using the sequencing-based neutralization assay (top) or a traditional one-virus versus one-serum fluorescence-based neutralization assay (bottom) (21,22). For the sequencing-based assay, the error bars represent the standard error of curves for replicate barcodes corresponding to the same strain from the same well; for the traditional neutralization assay the error bars represent the standard error of two replicate measurements calculated from separate wells. B) Correlation between NT50s for eight serum samples against three virus strains measured using the sequencing-based neutralization assay or a traditional fluorescence-based neutralization assay. Each point represents a different serum-virus pair (the virus strains shown are A/Hawaii/70/2019, A/Wisconsin/588/2019, A/England/220200318/2022), and points are colored by the day post-vaccination the sample was collected. The Pearson correlation (r) is indicated on the plot. C) Correlation between NT50s measured with sequencing-based neutralization assay versus HAI titers against the A/Hawaii/70/2019 virus strain for 30 serum samples collected on day 0 or day 30 post-vaccination. The dynamic range for the HAI titers was smaller than that used for neutralization assays (5-1280 vs. 20-393660).

### Neutralization of recent strains pre- and post-vaccination by individuals who did or did not receive a vaccine the previous year

We applied the sequencing-based neutralization assay to quantify the magnitude and specificity of the neutralizing H1 antibody response to influenza vaccination in samples from a randomized placebo-controlled trial of Flublok, a recombinant-HA influenza vaccine (26). We selected samples from individuals who were randomly allocated to receive influenza vaccination for the 2021-2022 winter (containing clade 5a.2 A/Wisconsin/588/2019 (H1N1) vaccine strain), stratified by whether they had or had not also received an influenza vaccine in the previous 2020-2021 season (containing clade 5a.1 A/Hawaii/70/2019 (H1N1) vaccine strain) (Table 1, Fig. 5). We selected 15 adult participants in each group (with or without a previous-year vaccination), ranging in age from 20-45 years at enrollment. For this study specifically, we chose individuals within each of the groups who had an HAI titer of <10 against the A/Hawaii/70/2019 (H1N1) vaccine strain prior to vaccination in year 1 (2020-2021); this criterion means that the study focuses on individuals with relatively low initial titers. None of the participants were infected with influenza during the study time frame because there was minimal influenza circulation at the study site in Hong Kong between the initial 2020-2021 blood draw and the 2021-2022 blood draws due to COVID-19 mitigation measures (27,28). We analyzed sera collected immediately before (day 0) and approximately 30 days after the participants were given the 2021-2022 vaccine (Fig. 5). In addition, for a subset of participants, we also analyzed sera collected approximately 182 days after vaccination (Table 1, Fig. 5).

**Figure 5.**
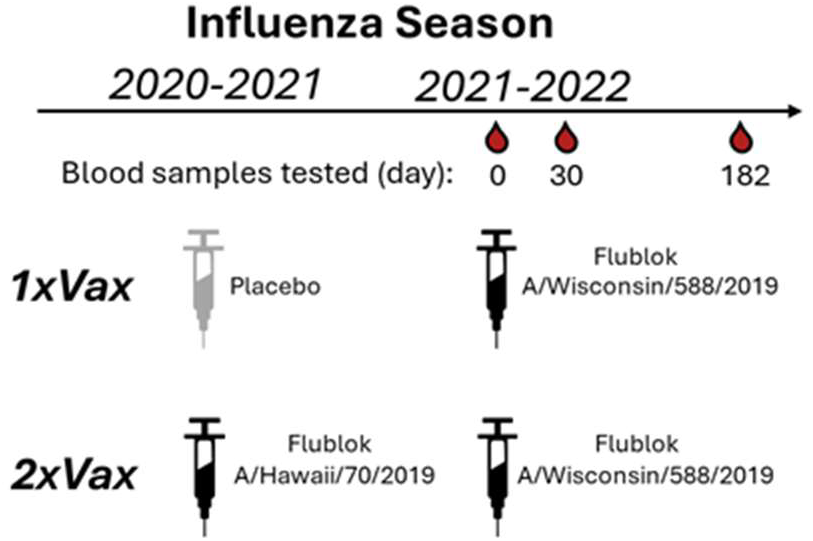
Schematic of study design. In the DRIVE study, a randomized vaccine trial of repeated annual vaccination of Flublok (a quadrivalent recombinant-HA vaccine), eligible healthy adults were randomly assigned at enrollment into five intervention groups of receiving a total of 1 to 5 annual Flublok vaccinations during 2020-2021 to 2024-2025 seasons, with blood specimens collected for analysis prior to receipt of the vaccine (day 0) as well as at day 30 and 182 post-vaccination in all participants. For the present analysis, we selected sera collected from study year 2 (2021-2022) from individuals pre-vaccination (day 0) and approximately 30 and 182 days post-vaccination for a subset of participants who had a pre-year 1 vaccination HAI titer <10 against the A/Hawaii/70/2019 (H1N1) vaccine strain, and who were assigned to either receive the placebo or the Flublok Vaccine (A/Hawaii/70/2019) in study year 1 (2020-2021 season). All selected participants received the Flublok Vaccine (A/Wisconsin/588/2019) in year 2 (2021-22 season), i.e. two groups of participants who were vaccinated in 2021-2022 and who had either prior year vaccination in 2020-2021 (“2xVax”) or not (“1xVax”) (Table 1).

**Table 1.** Description and demographics for participants selected for this study.

| **Study Group** | **Sex** | **Age (years)** | **Baseline HAI Titer** | **# with Day 0 and Day 30** | **# with Day 182** | **Vaccine received in 2020-2021** | **Vaccine received in 2021-2022** |
| --- | --- | --- | --- | --- | --- | --- | --- |
| *1x Vax* | Male | 24-44 | 5 | 8 | 6 | Placebo | FluBlok |
|  | Female | 20-29 | 5 | 7 | 4 | Placebo | FluBlok |
| *2x Vax* | Male | 23-45 | 5 | 8 | 7 | FluBlok | FluBlok |
|  | Female | 25-43 | 5 | 7 | 3 | FluBlok | FluBlok |

For each study participant, we generated neutralization landscapes that quantified the neutralization titers against all 36 H1 viral strains by the serum collected at each timepoint (see Fig. 6A for example landscapes, and Figures S4 and S5 for landscapes for all participants). These neutralization landscapes showed a variety of patterns. For instance, vaccination of participant D10066 induced a broad response against all viral strains (Fig. 6). Vaccination of participant D10011 induced a large increase in neutralizing titers against 5a.1 clade viruses, but a smaller increase in titer to most 5a.2 clade viruses other than the vaccine strain A/Wisconsin/588/2019 and a closely related strain (Fig. 6). Participant D10366, who had been vaccinated the previous season, had high pre-vaccination titers against 5a.1 clade viruses but no neutralizing response to vaccination in 2021-2022 (Fig. 6). A variety of other patterns can be seen by examining the neutralization landscapes for all participants in Fig. S4 and S5.

**Figure 6.**
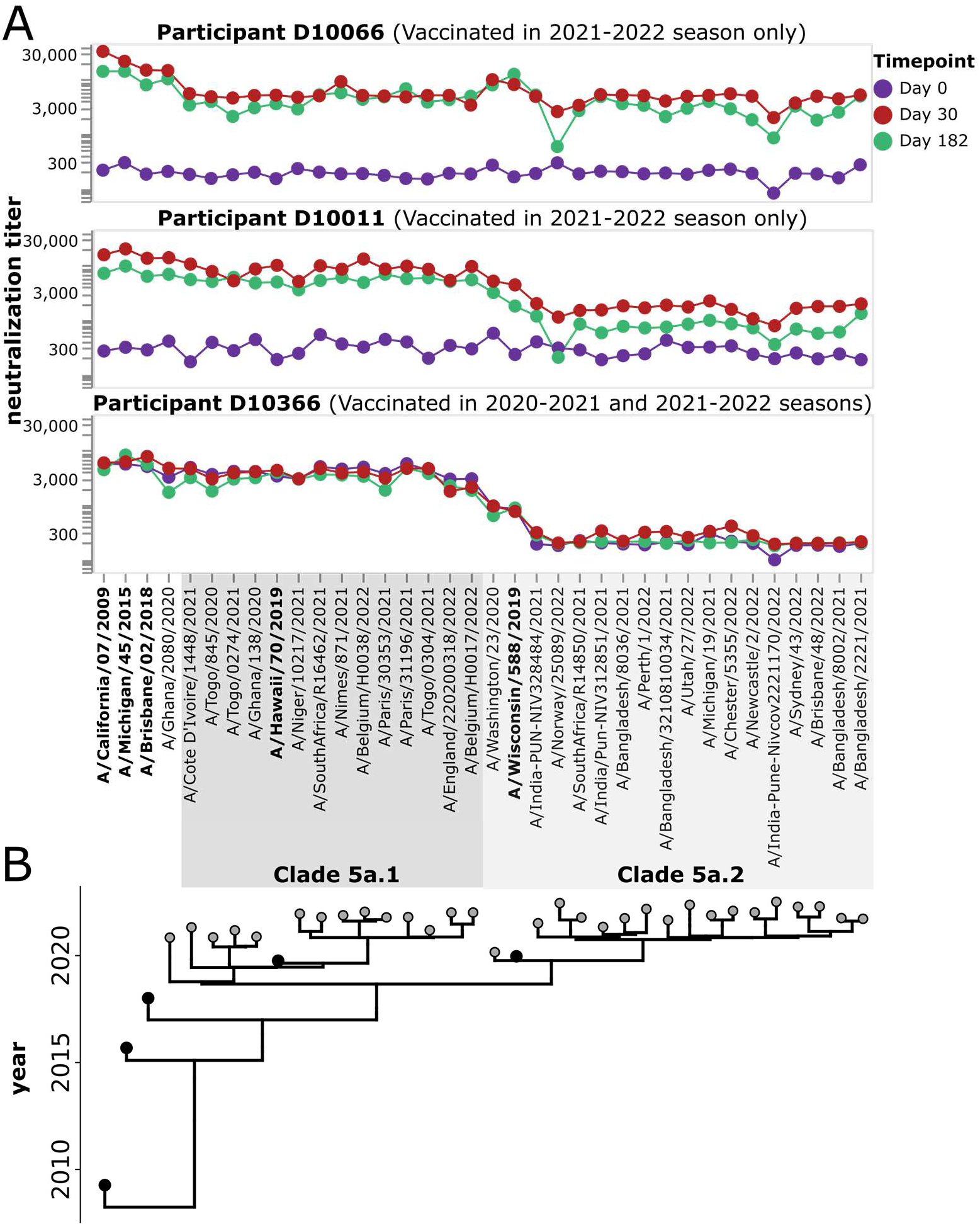
Example neutralization landscapes showing different strain- or clade-specific responses to vaccination. A) NT50s for three participants (D10066, D10011, and D10366) against all viral strains in the library for serum samples collected at 0, 30, or 182 days after receipt of the 2021-2022 vaccine. D10366 was also vaccinated in the previous season, while D10066 and D10011 were not vaccinated in the previous season. The participants responded differently to the 2021-2022 vaccination. Participant D10066 had a broad response to all viruses in the library. Participant D10011 had a stronger response to the 5a.1 clade than the 5a.2 clade, with the only strong response to a 5a.2 clade viruses being to the A/Wisconsin/588/2019 vaccine strain and a closely related strain. Participant D10366 did not respond to vaccination with increased titers to any strains but had high pre-existing titers to 5a.1 clade viruses probably due to vaccination in the previous season. See Fig. S4 and S5 for similar neutralization landscapes for all study participants. B). Phylogeny showing relationships among viral strains in the library. The names of the vaccine strains are in bold black text. Strains within clades 5a.1 and 5a.2 are indicated by shaded grey boxes.

Across the entire study cohort, participants who had been vaccinated the previous year had higher initial titers against recent viruses prior to receipt of the 2021-2022 vaccine compared to those who had not been vaccinated in previous year (Fig. 7A, S5). The pre-vaccination titers of these previous-year vaccinees were higher against 5a.1 clade than 5a.2 clade viruses, probably because the H1N1 component of the previous year (2020-2021) vaccine strain was a 5a.1 clade strain (A/Hawaii/70/2019). The group that received the placebo in 2020-2021 season had low pre-vaccination titers against all recent strains included in the library (Fig. 7A, S4), an expected result since study participants were chosen based on low initial baseline titers against the 2020-2021 vaccine strain.

**Figure 7.**
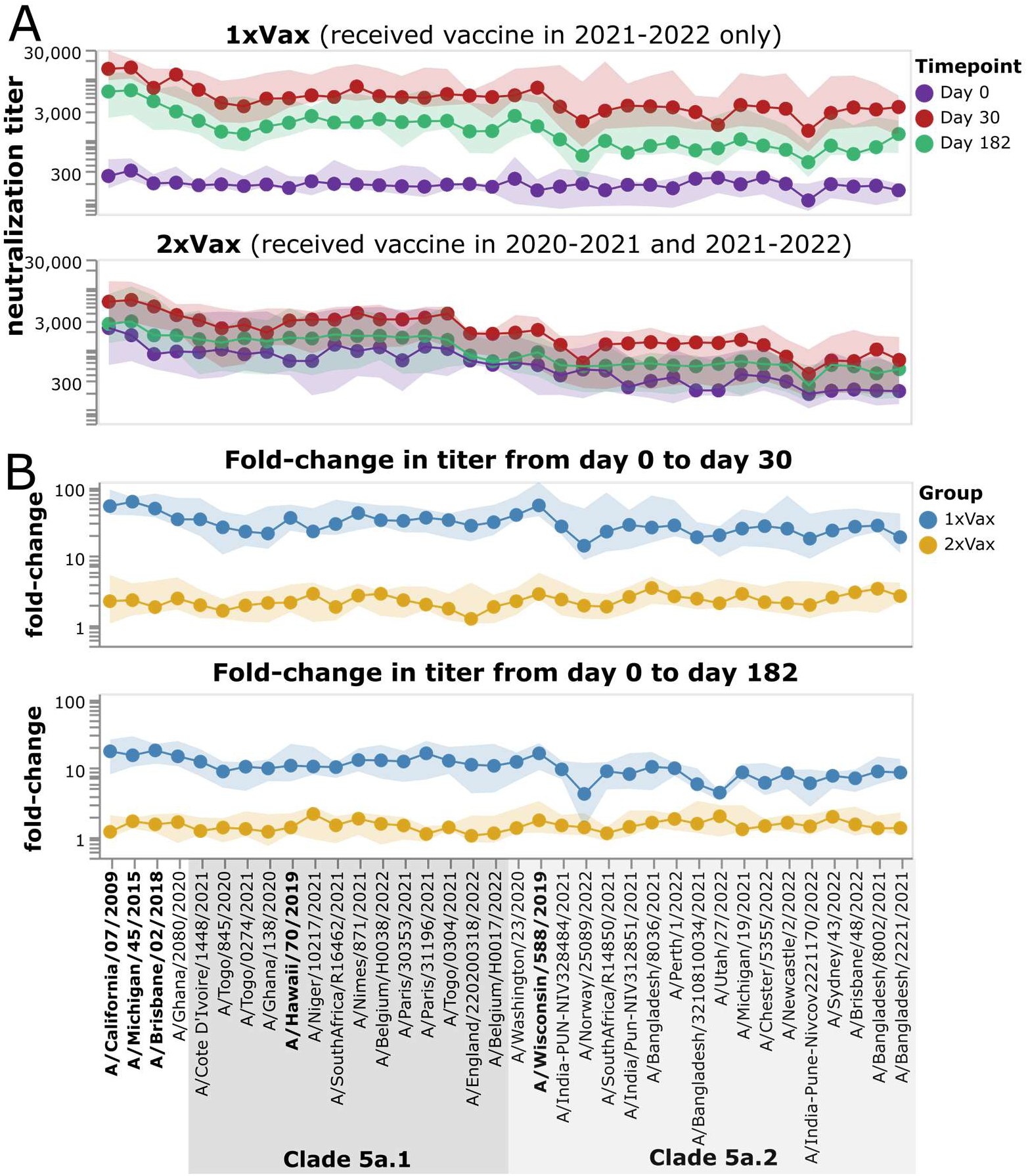
Impact of vaccination in 2021-2022 on neutralization titers across study participants who did or did not receive a vaccine the previous year. A) Neutralization titers against each virus strain at each timepoint for participants who were not vaccinated the previous year (top) or who also received a vaccine in 2020-2021. The points indicate the median titers across participants, and the shaded areas show the interquartile range. B) Fold change in titer at day 30 or day 182 post-vaccination relative to the day 0 titer. Points indicate the median fold change across participants, and the shaded areas show the interquartile range. Strains within clades 5a.1 and 5a.2 are indicated by shaded grey boxes. The names of the vaccine strains are in bold black text. This figure shows only participants who had serum samples from all three timepoints (days 0, 30, and 182 post vaccination); see Fig. S6 for plots that include participants who only had samples from the day 0 and 30 timepoints.

Following vaccination in 2021-2022, most individuals who had not been vaccinated in the previous year responded with large increases in neutralizing antibody titers against all strains (Fig. 7, S5). In contrast, only some individuals who had also been vaccinated in the previous year showed an appreciable response to vaccination in 2021-2022, and the magnitude of this response was generally smaller than in the group with a previous-year vaccination (Fig. 7, S4, S5).

Specifically, only 8 of 15 participants in the previous-year vaccination group had at least a four-fold increase in titer against the vaccine strain at 30-days post vaccination, while 14 of 15 individuals in the group without a previous-year vaccination had at least a four-fold increase (Fig. S4, S5). Both groups showed some decrease in titer from day 30 to day 182, consistent with other studies showing waning of the antibody response to influenza vaccination within a few months (29,30).

However, despite the much stronger response to vaccination in the first-time versus previous-year vaccinees, the absolute neutralization titers after vaccination are not dramatically different between the two groups (Fig. 8, S6). At 30 days post-vaccination, the first-time vaccinees do have higher titers to nearly all strains than the previous-year vaccinees (as assessed by the median titers across the groups), with some of the differences being statistically significant—but there is substantial overlap between the two groups, and the differences in absolute titers are much more modest than the differences in fold change (Fig. 8, S6). Many of the strains for which the first-time vaccinees have higher absolute titers than the previous-year vaccinees are in the 5a.2 clade, which is the same clade as the 2021-2022 vaccine strain, A/Wisconsin/588/2019 (Fig. 8, S6). However, the modest differences in absolute neutralization titers between the two groups at 30 days post-vaccination are nearly gone by 182 days post-vaccination (Fig. 8). Overall, these results show that while the titer rise caused by vaccination is much larger in participants who have not been vaccinated the previous year compared to those who received a previous-year vaccine, the actual titers post-vaccination are fairly similar, with first-time vaccinees having slightly higher titers at 30 days, but not 182 days post vaccination.

**Figure 8.**
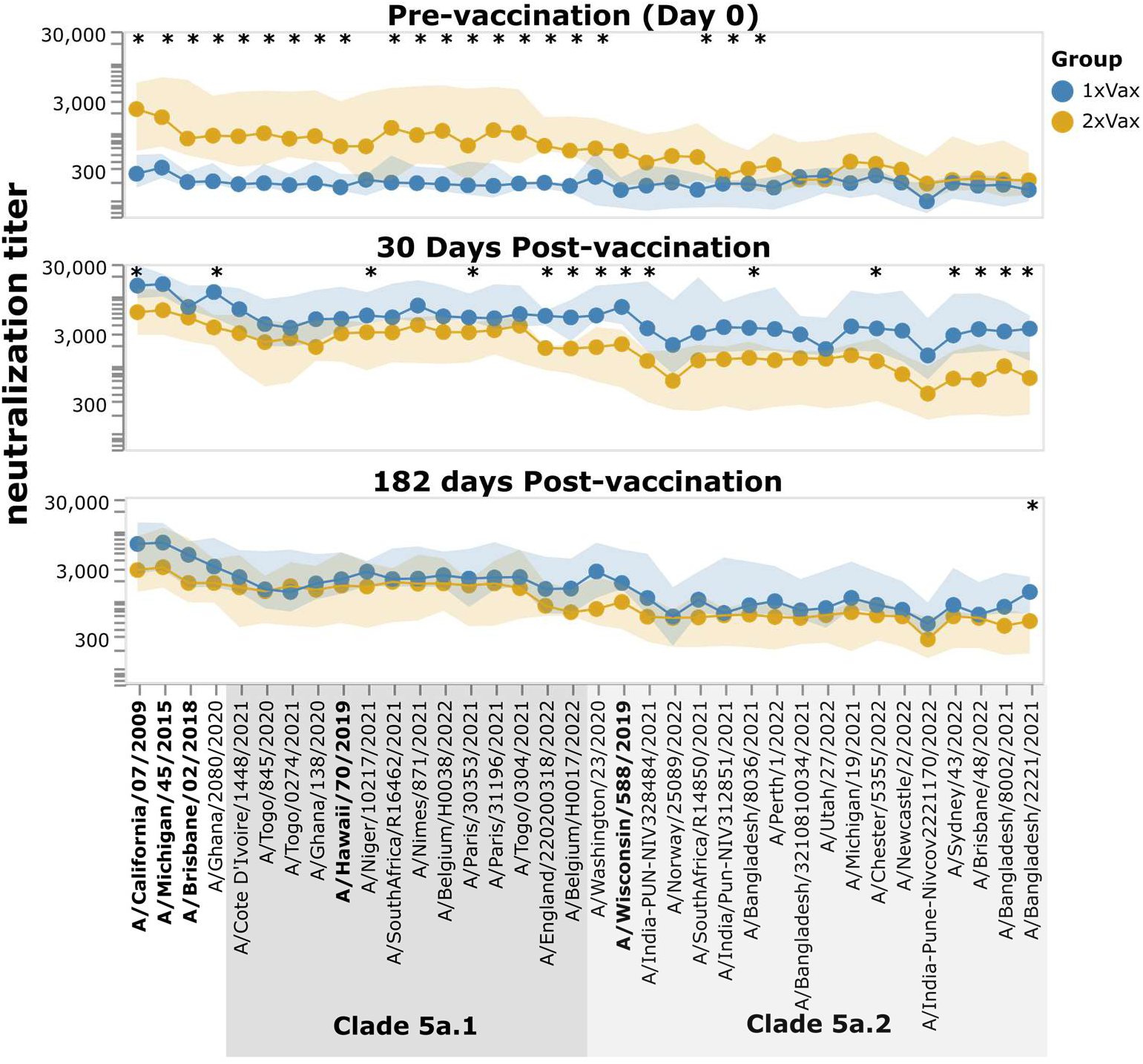
Neutralization titers after receipt of 2021-2022 vaccine among participants that did or did not also receive a vaccine in the previous year. The points show the median neutralization titer across study participants in that group, and the shaded areas show the interquartile range. The names of the vaccine strains are in bold black text. Strains within clades 5a.1 and 5a.2 are indicated by shaded grey boxes. Strains with a significant difference in median titer between groups as assessed by a Mann-Whitney U Test are indicated with an asterisk at the top of each plot panel. Titers against all strains for each individual study participant are shown in Fig. S4 and S5. This figure shows only participants who had serum samples from all three timepoints (days 0, 30, and 182 post vaccination); see Fig. S6 for plots that include participants who only had samples from the day 0 and 30 timepoints.

### Identification of specific viral strains with reduced neutralization by some study participants

The fact that we generate full neutralization landscapes for each study participant against many viral strains enables identification of specific viruses with reduced neutralization by specific participants. For example, several individuals have markedly lower post-vaccination neutralization titers against a small subclade of the 5a.1 clade (represented by viruses A/Belgium/H0017/2022 and A/England/220200318/2022) than against other viral strains. A subset of individuals with limited neutralization against this particular subclade is shown in Fig. 9. These more poorly neutralized viral strains carry HA mutations P137S and G155E, both of which are in known antigenic regions of the H1 HA (the Ca and Sb epitopes, respectively) (15). Only one other strain in the library has either of these mutations: P137S is found in the 5a.2 strain A/Norway/25089/2022, but that strain does not have reduced neutralization by any of these participants (Fig. 9). This fact suggests that the reduced neutralization of the subclade containing A/Belgium/H0017/2022 and A/England/220200318/2022 could be due to G155E, although it also could be due to the combined action of several mutations in a fashion that depends on viral genetic background.

**Figure 9.**
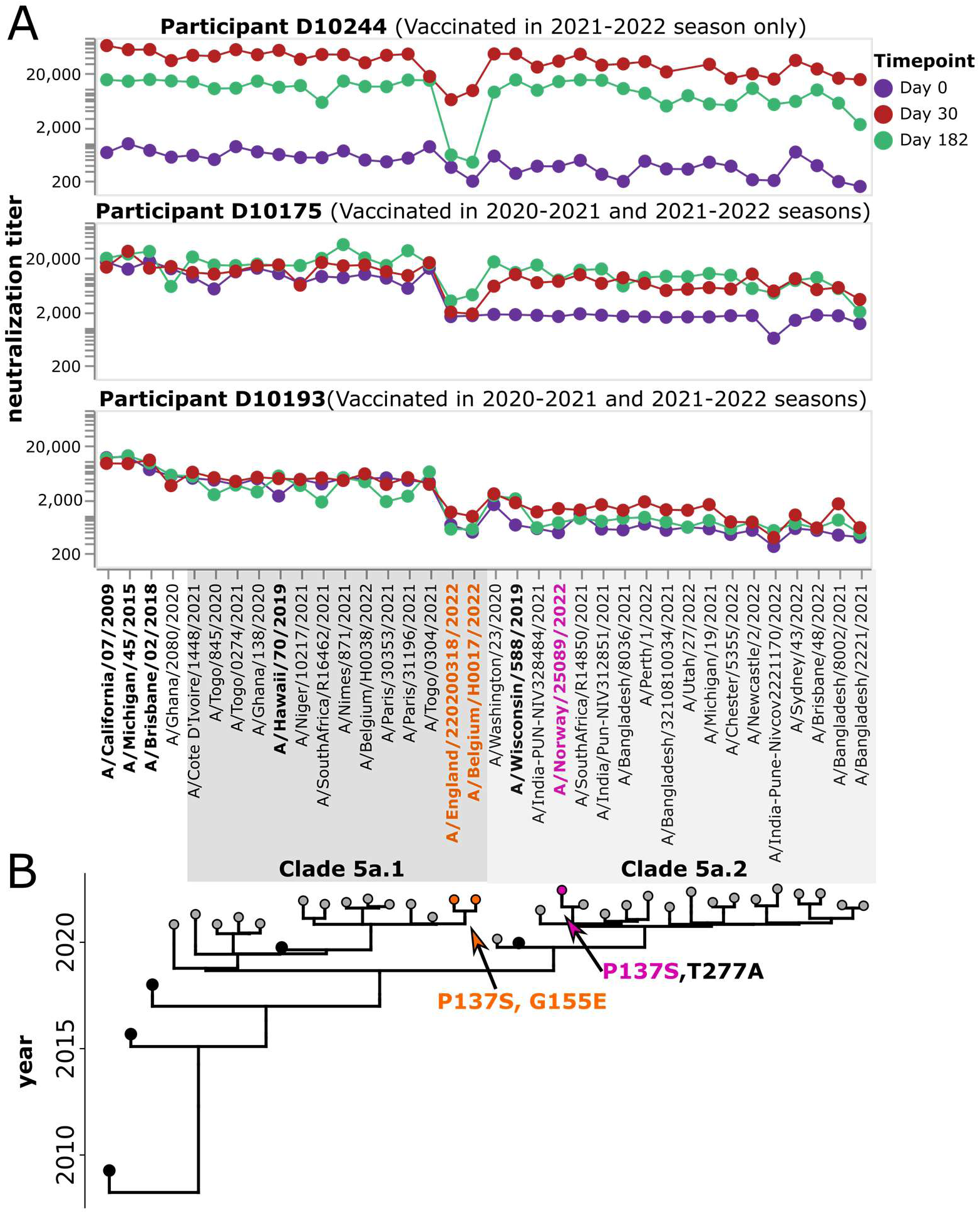
Some participants have low neutralizing titers to viruses in a specific subclade of 5a.1. A) Neutralization landscapes for three study participants with low titers to a subclade of 5a.1 viruses consisting of A/Belgium H0017/2022 and A/England/220200318/2022. B) Phylogeny of the viral strains included in the library, with orange indicating strains that contain the HA mutations P137S and G155E, and pink indicating a strain that contains P137S but not G155E.

Inspection of Fig. S4 and S5 also reveals other instances where specific participants have enhanced or reduced neutralization of specific viral strains. Overall, these observations emphasize how neutralization titers are both participant- and virus-specific.

## Discussion

We describe a new sequencing-based neutralization assay that enables measuring titers against many influenza viruses with the same serum volume and a similar workflow as a traditional single-virus neutralization assay. In the current study, we applied the assay to simultaneously measure titers to viruses with 36 different H1N1 HAs, but the same workflow could be applied to influenza A viruses of any HA subtype, or a library of viruses with HAs from different subtypes. We estimate that the method could be extended to simultaneously measure titers for as many as ∼100 different viruses at once, with the upper bound limited by the fact that the library needs to be small enough that each well in a 96-well plate has many cells infected by each viral barcode while maintaining only a modest multiplicity of infection (MOI). Like traditional neutralization assays, this sequencing-based assay is conducted in 96-well plates, which can facilitate the processing of many serum samples at once.

The ability to measure neutralization titers against many viral strains is important because influenza HA is so antigenically variable. Humans typically have high neutralizing titers to historical strains to which they have been exposed but are at risk of re-infection from new strains with HA mutations that erode this neutralization. Traditional neutralization or HAI assays can measure titers against a handful of strains (21,31–34), whereas approaches like deep mutational scanning (35) can measure the effects of all individual HA mutations in the genetic background of a specific strain. While various technical improvements have been made to increase the throughput of one-virus versus one-serum neutralization assays (36,37), the effort required to perform these assays still scales linearly with the number of viral strains being tested. In contrast, the new sequencing-based neutralization assay we have developed can be applied to ∼100 viral strains just as easily as it can be applied to a single virus strain. We anticipate the assay can be applied for several purposes, including assessing heterogeneity in the specificity of human immunity to different viral strains, providing antigenic data that can inform forecasting models for influenza vaccine-strain selection (38,39), and characterizing the breadth of responses induced by different vaccine regimens.

Here we applied the sequencing-based neutralization assay to examine how repeat vaccination influenced neutralizing titers against recent H1N1 strains that circulated in 2020-2022.

Consistent with the antibody ceiling effect, in which individuals with higher initial titers typically show smaller increases in titer following vaccination (40–42), the increase in neutralizing titer following vaccination was higher in individuals who had not been vaccinated the previous year compared to those who did receive the previous-year vaccine. The extent of this difference in fold-change between the two groups in our study may be magnified by our choice to limit our analysis to individuals who had low titers prior to receiving the vaccine or placebo in the first year. Still, despite the lower fold-increase in individuals with a previous-year vaccination, the absolute neutralization titers were fairly similar between the two groups because the previous-year vaccinees started with higher titers. There was a slight trend for the first-time vaccinees to have higher titers than the previous-year vaccinees at 30 days post-vaccination, but by 182 days post-vaccination the differences had mostly disappeared. Importantly, there was also substantial heterogeneity both among individual study participants and across viral strains. For instance, we identified specific subclades of viruses that were poorly neutralized by some study participants. This heterogeneity emphasizes the value of a high-throughput assay that can measure the neutralization titers of many different human sera against many different viruses, and may help explain why vaccination fails to protect against infection in some individuals.

Using the sequencing-based neutralization assay, with a barcoded library in hand, one researcher was able to collect 2,880 neutralization titers in ∼4 weeks. Broader implementation of this assay could greatly expand the density of information that can be collected on the ability of different individuals to neutralize different influenza virus strains—an advance that would have multiple important applications. For instance, the rapid identification of antigenically distinct strains could aid in vaccine strain selection (43,44). Similarly, the ability to assess titers against many strains could help identify vaccination regimens that induce broader immunity, as well as help with understanding current immune heterogeneity in the human population due to variation in exposure history or other factors(18,45–47). To facilitate the wide use of sequencing-based neutralization assays, we provide a detailed protocol both in the methods section of this paper and on protocols.io (DOI: https://dx.doi.org/10.17504/protocols.io.kqdg3xdmpg25/v1 ).We have also made available a computational pipeline suitable for analyzing and quality-controlling the thousands of neutralization curves that can be generated using sequencing-based assays (https://github.com/jbloomlab/seqneut-pipeline).

## Acknowledgements

This work was supported in part by the NIH/NIAID award R01AI165821 to TB and JDB, U01AI153700 to SC and BJC, and contract 75N93021C00015 to JDB, SC, BJC, and TB. JDB and TB are Investigators of the Howard Hughes Medical Institute. This research was supported by the Genomics & Bioinformatics Shared Resource (RRID:SCR_022606) of the Fred Hutch/University of Washington/Seattle Children’s Cancer Consortium (P30CA015704), and by Fred Hutch Scientific Computing (NIH grants S10-OD-020069 and S10-OD-028685). BJC is supported by the Theme-based Research Scheme (Project No. T11-712/19-N) of the Research Grants Council of the Hong Kong Special Administrative Region, China. We thank Richard Webby at St. Jude’s Children’s Research Hospital for providing virus strains for HAI assays, and Lewis Siu for assistance in virus preparation for these assays. We thank Caroline Kikawa for helpful feedback. We gratefully acknowledge all data contributors for the sequences downloaded from GISAID, including the authors and their originating laboratories responsible for obtaining the specimens, and their submitting laboratories for generating the genetic sequence and metadata and sharing via the GISAID Initiative, on which part of this research is based. A list of laboratories contributing sequences for strains used in this work is provided in https://github.com/jbloomlab/flu_seqneut_DRIVE_2021-22_repeat_vax/tree/main/librarydesign.

## Author Contributions

Conceptualization, ANL and JDB. Methodology, ANL and JDB. Investigation, ANL, RALT, JH. Software, ANL, JDB, and WH. Validation, ANL, RALT, LT, and SSW. Resources, SMSC, NHLL, SC, BJC. Writing – Original Draft Preparation – ANL and JDB. Writing – Reviewing and editing, all authors. Funding acquisition, JDB, TB, SC, BJC.

## Materials and Methods

### Data and code availability

See https://github.com/jbloomlab/flu_seqneut_DRIVE_2021-22_repeat_vax for all data and computer code. Specifically:

- The neutralization titers (NT50s) for each virus-serum pair are at https://github.com/jbloomlab/flu_seqneut_DRIVE_2021-22_repeat_vax/blob/main/results/aggregated_titers/titers.csv
- Details about the samples are at https://github.com/jbloomlab/flu_seqneut_DRIVE_2021-22_repeat_vax/blob/main/data/sample_metadata_forplots.csv
- The barcodes assigned to each viral HA are at https://github.com/jbloomlab/flu_seqneut_DRIVE_2021-22_repeat_vax/blob/main/data/viral_libraries/pdmH1N1_lib2023_loes.csv
- The barcodes for the neutralization standards are at https://github.com/jbloomlab/flu_seqneut_DRIVE_2021-22_repeat_vax/blob/main/data/neut_standard_sets/loes2023_neut_standards.csv
- Maps of all plasmids including the HA sequences are at https://github.com/jbloomlab/flu_seqneut_DRIVE_2021-22_repeat_vax/tree/main/plasmids
- Plots of neutralization curves and other key results are at https://jbloomlab.github.io/flu_seqneut_DRIVE_2021-22_repeat_vax/
- Other results files, including all barcode counts and the fraction infectivities used to fit all neutralization curves, are in the “results” subdirectory at https://github.com/jbloomlab/flu_seqneut_DRIVE_2021-22_repeat_vax

### Design of barcoded HAs

Influenza is a segmented, negative sense RNA virus. Proper packaging of the viral RNA (vRNA) segments into the virion relies on segment-specific RNA sequences called “packaging signals” which are present at the 3’ and 5’ ends of each segment and include both coding and noncoding regions (48,49) (Fig S1A). To insert a unique barcode into the HA genomic segment without substantially disrupting vRNA packaging, we designed a chimeric construct with a duplicated packaging signal at the 5’ end of the negative-sense HA vRNA (Fig. S1B) (13,14,50–54), see plasmid maps in https://github.com/jbloomlab/flu_seqneut_DRIVE_2021-22_repeat_vax/tree/main/plasmids. As all non-HA genes used to generate the barcoded influenza viruses are from the lab-adapted A/WSN/1933 (H1N1) strain, we used terminal packaging signals from this strain to better facilitate incorporation of HAs from various recent H1N1 strains into the virions (Fig S1B) (53–55). At the 3’ end of the negative-sense HA vRNA, the native 3’ packaging signal (defined as the 32-nucleotide noncoding region and first 67 nucleotides of the coding sequence of HA (48)) from A/WSN/1933 HA was used (Fig S1B). We then incorporated the ectodomain from the HA strain of interest, spanning amino acids 23-520. The transmembrane domain and cytoplasmic tail of the HA, spanning the last 46 amino acids, are replaced with the homologous 46 amino-acid region from the WSN strain followed by a pair of consecutive stop codons (144 nucleotides in total). We incorporated synonymous mutations into the last 105 nucleotides of this region encoding the C-terminal portion of the HA to remove the packaging signals without altering the encoded protein (Fig S1B). This synonymous re-coding reduces homology between the duplicated region within the coding sequence and the terminal packaging sequence used for incorporating the segment into the virion, which may be important for reducing intra-segment recombination. Immediately following the consecutive stop codons, we incorporated a random 16-nucleotide barcode followed by the Illumina read 1 priming sequence. We then included the last 150-nucleotides of the negative sense 5’ HA vRNA from the WSN strain (this is the full WSN HA 5’ packaging signal). A stop codon was introduced into what would have been the coding sequence of this final region with the goal of improving barcode retention: if recombination between the partially homologous regions immediately upstream and downstream of the barcodes occur, this stop codon should truncate the HA protein therefore yield a non-functional virus (Fig S1B, and https://github.com/jbloomlab/flu_seqneut_DRIVE_2021-22_repeat_vax/tree/main/plasmids).

### Selection of recent human H1N1 HAs for includes in the library

To select strains for inclusion in barcoded influenza library, we first identified emerging clades on using a Nextstrain (56,57) build for human H1N1pdm sequences from October 12, 2022 (https://github.com/jbloomlab/flu_seqneut_DRIVE_2021-22_repeat_vax/tree/main/librarydesign). This was done using a method available through augur on Nextstrain (similar to https://github.com/nextstrain/augur/blob/master/scripts/identify_emerging_clades.py). Clades were selected based on the following criteria: they must contain at least one HA1 amino acid mutation compared to the parent clade and have descendants that were represented at a global frequency of greater than 1%. This yielded a set of 33 clades; however, two clades were excluded. One of the clades was eliminated as all members of that clade overlapped with an included subclade, and the other was excluded as all sequences were from 2020 or before, indicating that this clade was likely no longer circulating. Next, we manually selected a sequence from each of these clades that contained the clade-defining mutations and was close to the node of the clade, such that the selected strains did not include additional mutations which were not shared with all members of the clade. This yielded 31 strains that represented the diversity of viruses that were circulating in the fall of 2022. This list of strains is at https://github.com/jbloomlab/flu_seqneut_DRIVE_2021-22_repeat_vax/tree/main/librarydesign. In addition to the selected recent strains, the following vaccine strains were also included in the library: A/California/07/2009, A/Michigan/45/2015, A/Brisbane/02/2018, A/Hawaii/70/2019, and A/Wisconsin/588/2019.

### Generation of barcoded influenza viruses

The ectodomains of the hemagglutinin proteins from selected influenza virus strains were ordered as synthetic DNA fragments from Twist Biosciences and cloned with a barcoded fragment encoding the last 46 amino acids of WSN HA as 3-segment assembly reaction into a construct containing the 3’ non-coding region of the packaging signal and signal peptide from A/WSN/1933 influenza HA and the a Read 1 Illumina sequence and the full 5’ packaging signal from A/WSN/1933 virus using Hifi Assembly Mastermix (NEB). The backbone for this cloning reaction was a pHH21 plasmid (16) which had been modified to contain the packaging sequences of WSN HA, with restriction enzyme cut sites flanking the 3’ packaging signal, specifically, a *BamHI* site, which results in cleavage after nucleotide 67 of the viral HA genomic segment, and an *XbaI* immediately preceding the read 1 sequence to allow for incorporation of unique barcodes into each plasmid. The full plasmid map for the vector used for this reaction made available as *2805_WSNHAflank_GFP_NterminalWSNHA_duppac-single-stop.gb* in https://github.com/jbloomlab/flu_seqneut_DRIVE_2021-22_repeat_vax/tree/main/plasmids. Each strain was cloned with 3-4 barcodes/strain, and these were sequence-verified with whole plasmid sequencing through Primordium Labs. See https://github.com/jbloomlab/flu_seqneut_DRIVE_2021-22_repeat_vax/tree/main/plasmids for the Genbank maps for all plasmids.

To generate actual viruses from these plasmids, the unidirectional reverse genetics plasmids encoding all barcoded variants of the same HA sequence were pooled at equal concentrations and used to generate influenza viruses containing unique barcodes by reverse genetics, with the WSN pHW18* series of plasmids for all seven non-HA viral genes (17), and pHAGE2-EF1aInt-TMPRSS2-IRES-mCherry-W(58). Briefly, 5e5 293T cells and 5e4 MDCK-SIAT1-TMPRSS2 cells(58) were plated in a 6-well dish in D10 media (DMEM supplemented with 10% heat-inactivated fetal bovine serum, 2 mM L-glutamine, 100 U per mL penicillin, and 100 µg per mL streptomycin). Approximately 24 hours after plating cells, a master mix containing 240 ng/well of each of the reverse-genetics plasmids for WSN virus and 80 ng of the TMPRSS2 containing plasmid was prepared in 100 uL DMEM with 3 uL BioT reagent. This was incubated for 15 minutes at room temperature and then added dropwise to the plate.

Approximately 20 hours post-transfection, media was removed, cells were washed once with PBS (phosphate-buffered saline) and 2 mL Influenza Growth Media (Opti-MEM supplemented with 0.1% heat-inactivated FBS, 0.3% bovine serum albumin, 100 µg per mL of calcium chloride, 100 U per mL penicillin, and 100 µg per mL streptomycin) was added to cells. At ∼65 hours post-transfection cell supernatants containing the barcoded influenza viruses were collected and centrifuged for 4 min at 845 xg to remove cell debris. Aliquots of clarified viral supernatant were then frozen at -80°C for storage. To expand viruses to high titer, all virus pools generated by reverse genetics were passaged once in MDCK-SIAT1-TMPRSS2 cells (58). For this, MDCK-SIAT1-TMPRSS2 cells were seeded at a density of 4e5 cells/well in a 6-well dish in D10, and at 4 hours after seeding were washed with PBS and 2 ml of Influenza Growth Medium was added to each well,, Each well was then inoculated with 200 uL of the virus supernatant from reverse genetics and the virus was allowed to grow in the cells for ∼40 hours. Cell supernatants were then collected and clarified by centrifugation at 845 xg for 4 min, and aliquots of clarified viral supernatants were stored at -80°C for use in the sequencing-based neutralization assay.

### Pooling of influenza virus library strains

To generate a pooled library where each strain was represented at approximately equal transcriptional titers, we used a sequencing-based titering method. First, an equal volume mix of all of the passaged barcoded virus strains that were to be incorporated into the virus library was made and titered. The TCID50 for this initial pool was ∼35,900 TCID50/uL on MDCK-SIAT1-TMPRSS2 cells. Serial dilutions of this pool were used to inoculate a single well of a 96-well plate containing 50,000 MDCK-SIAT1 cells, and after 16 hours the viral barcode frequencies were determined by deep sequencing as described in a later section. We determined the relative fraction of all barcode counts that corresponded to each strain (Fig S2A, left panel). We then took the fraction of the library that each strain should represent in an ideal equal-representation pool (the reciprocal of the number of strains, in this case 1 / 36) and divided this number by the fraction in the initial equal-volume pool to find the ratio at which that strain should be added. We multiplied this ratio by 200 uL to determine the volume to add for each strain to generate the final pool, with sufficient volume to run ∼100 neutralization assay plates. The resulting pool was titered by TCID50 and found to be at ∼39600 TCID50/uL. We sequenced cells infected with the transcriptional-titer balanced pool and found that all strains were within approximately 2-fold representation of each other (Fig. S2A, right panel). We also examined the relative representation of each barcode for each strain (Fig. S2B). There is variability in barcode representation for each strain (Fig. S2B), however, as the replicate barcodes for each strain were rescued by reverse genetics as a pool, these ratios cannot be manipulated at this stage. Independent rescues could be performed for each barcoded strain to be able to balance barcodes, rather than strains, but that would substantially increase the number of rescues needed to prepare a barcoded library. The yield of virus library obtained with the approach described above depends on the titers of the viruses included in the library. If viruses are generated at titers between 1,000 TCID50/uL – 10,000 TCID50/uL, with 2 mL of each strain per strain included in the library, and a similar library size (36 strains), one should have sufficient volume to run between 26-260 sequencing-based neutralization assay plates, or sufficient volume to perform measurements against the library for ∼800-8000 serum samples. However, the relevant titer is the transcriptional titer of the library (how many virions can infect cells to transcribe the HA barcode), not TCID50/uL. This transcriptional titer should be measured experimentally for each new library as described in a later section. The necessity of this rebalancing depends on how different the titers are between the different strains one wishes to include in the library. For this library, only a small number of strains were poorly represented in the equal volume pool, however if there is significant variability in growth of library strains, this rebalancing would be essential to ensure that each strain is adequately represented in the final pool.

### Verifying barcode stability in the HA genomic segment

Our lab has previously confirmed that barcodes introduced into the influenza HA gene segment using a broadly similar strategy to that in this paper are stably retained by the virus (13,14). But to confirm that the barcode region was genetically stable within the exact HA constructs in the current study, we analyzed the HA sequences of the virus library generated by pooling the passaged viral stocks. For this analysis, RNA was isolated from the viral supernatant of the pooled library using the RNeasy PLUS Mini Kit (Qiagen, 74104). Lysis buffer (RLT) was mixed with the viral supernatant sample and 70% ethanol was added, the sample was then applied to the RNAeasy purification column and RNA was isolated according to the protocol supplied by the manufacturer. The RNA for each sample was eluted in 22 uL of RNase-free water. Reverse transcription was performed using 5 uL of the extracted RNA according to manufacturer’s protocol for the SuperScript III First-Strand Synthesis SuperMix kit (Thermo Scientific), with a gene-specific primer *full-length_HA_Fwd:* 5’-*AGCAAAAGCAGGGGAAAATAAAAACAACC*-‘3. Following cDNA synthesis, PCR using *full-length_HA_Fwd* primer and *full-length_HA_Rev:* 5’-*AGTAGAAACAAGGGTGTTTTTCCTTATATTTCTGAAATCC*-3’ was performed to amplify the HA segment. A plasmid containing barcoded HA was used as a control to verify expected size. We examined both a single virus strain after initial reverse genetics and blind passage and found that the full-length barcoded construct was maintained. In the pooled library, some deletions are observed in the HA segment (Fig. S7A). We Sanger-sequenced all bands observed in the gel and verified that all of them included the barcode region, the largest band is the intact, full-length HA segment. Significant diversity in nucleotide identity was observed within the barcode region of this segment, as expected. The next largest band contains a deletion within the HA ectodomain. This segment also had diversity barcodes associated with it, indicating that this deletion likely occurred in multiple barcoded HAs. Finally, the smallest segment is a deletion product missing much of the HA protein sequence, but containing only a single barcode, indicating that it likely originated in one of the passages from a single strain. This barcode is not associated with any of the sequence confirmed HAs included in the library, so no curves were generated for this sequence. The presence of some amount of internal deletions in influenza gene segments including HA is common (59), and does not pose any problem for our experimental approach as long as the barcode region is retained (which is the case here). To confirm that internal deletions were not rampant in any other influenza genomic segments (which would suggest high amounts of defective interfering particles), we performed cDNA synthesis and PCR as described above using primers that annealed to all 8 segments of the influenza genome as previously described by others (60). All segments appear to be present in these stocks, suggesting that they are not dominated by defective interfering particles (Fig. S7B).

### Generation of RNA spike-in control

Our approach used an RNA spike-in control to convert sequencing counts to an absolute fraction of viral infectivity remaining (Fig. 1). To generate a spike-in control single-stranded RNA that contained known barcodes and resembled the vRNA of barcoded influenza viruses, we first cloned the protein sequence for green fluorescent protein in place of the HA coding sequence in the plasmid backbone we used for the barcoded HAs (see schematic in Fig. S1C and the full plasmid map 3808_*WSNHAflank_GFP_WSNtermini_definedbarcodes_forSpikeinControl*, available in https://github.com/jbloomlab/flu_seqneut_DRIVE_2021-22_repeat_vax/tree/main/plasmids), and then sequence-verified 10 plasmids containing unique barcodes with whole plasmid sequencing. We then generated a linear PCR template from the pooled set of barcoded plasmids using primers complementary to the termini of the vRNA encoded in the plasmid, *full-length_HA_Fwd:* 5’-*AGCAAAAGCAGGGGAAAATAAAAACAACC*-3’ and *full-length_HA_Rev:* 5’-*AGTAGAAACAAGGGTGTTTTTCCTTATATTTCTGAAATCC*-3’ respectively. The PCR product was run on a 1% agarose gel at 85 V for 35 min and gel extracted using the Nucleospin Gel Extraction Kit (Takara). We then used PCR with KOD Polymerase 2x Mastermix (Millipore Sigma) to append a T7 promoter sequence, using the following primers: *Add_T7_Rev:* 5’-*TTACGATAATACGACTCACTATAGGGAGTAGAAACAAGGGTGTTTTTCCTTATATTTCTG*-3’ and *full-length_HA_Fwd:* 5’-*AGCAAAAGCAGGGGAAAATAAAAACAACC*-‘3, with 500 ng of the purified linear template. Then we used T7 Ribomax Express (NEB) *in vitro* RNA synthesis kit to generate single-stranded negative sense RNA that resembles the vRNA molecule. To remove DNA template after *in vitro* transcription, 0.5 uL of RQ DNAse was added to the reaction and incubated another 15 min at 37°C. The Monarch RNA Cleanup Kit (NEB) was used to purify the resulting RNA. The product was quantified by Nanodrop, diluted to 200 pM, aliquoted for use in each experiment, and stored at -80°C. We verified that the RNA spike-in stock was free of DNA template by performing cDNA synthesis using the iScript Select cDNA Synthesis Kit, with different concentrations of template (0.12 - 12 ng) with and without the reverse-transcriptase enzyme. PCR amplification was run with the Round 1 PCR primers and the product was run on a 1% agarose gel at 85 V for 1 hr. No product was observed in the no-template and no-enzyme controls, indicating that spike-in was free of contaminating DNA.

### Transcriptional titering to determine the amount of virus library to add to each well so that HA viral barcode counts can be accurately converted to fraction viral infectivity

The sequencing-based neutralization assays require that the number of sequencing counts for each HA viral barcode normalized by the spike-in RNA counts be directly proportional to the number of virions encoding that barcode that infected cells (Fig. 1). This condition will be met as long as the multiplicity of infection (MOI) is not too high, since at low to modest MOIs an increased number of virions infecting cells will produce more barcoded viral RNA, but at very high MOIs the capacity of infected cells to produce viral RNA becomes saturated. To identify the amount of virus library that can be added to a well while maintaining linearity between the amount of input library and the resulting spike-in normalized HA counts, we performed dilution series where wells were infected with different amounts of virus library and then sequenced as described above using 50,000 MDCK-SIAT1 cells/well with serial dilutions of the virus library at known TCID50s. As described above, we added a constant amount of the RNA spike-in control (50 uL of 2 pM RNA) in lysis buffer to each well containing cells infected with the virus pool. We then sequenced the barcodes in the well and identified the highest amount of virus library we could add and still be in the range where a two-fold decrease in input virus library led to a two-fold decrease in the counts of viral barcodes relative to spike-in control barcodes. We found that adding 25,000 TCID50/well (or a volume of ∼0.6 uL/well of library) and 50,000 MDCK-SIAT1 cells per well was appropriate for this particular library; note however that the TCID50 to cell ratio could vary for other libraries depending on the ratio of TCID50 to transcriptionally active virions. Our library has a total of 110 barcoded variants (the 36 different HAs each have 2-4 barcodes), corresponding to ∼220 TCID50 per well when using the amount of virus chosen for our experiments. These values present some anticipated limits to the library size that might be able to be characterized by the sequencing-based neutralization assays. At a coverage of at least 100 TCID50/well, and a maximum inoculum of 25,000 TCID50/well, with 3 barcodes per variant, we anticipate that up to 80 variants per well could be characterized with this method. A smaller number of barcodes per variant might be employed to increase the library size. The experiments described in this subsection should be repeated for each new library, as different stocks of virus may have different numbers of transcriptionally active particles.

### Sequencing of HA barcodes in virally infected cells

All viral infections for barcode sequencing were performed using MDCK-SIAT1 cells. We used these cells rather than the MDCK-SIAT1-TMPRSS2 cells to limit secondary viral replication (TMPRSS2 cleaves HA in producing cells to activate it for infection) (61–63). The cells were plated with influenza virus library (at indicated concentrations) with 50,000 cells per well in 96-well plates in IGM in a total volume of 150 uL. At 16 hours post-infection, cells were washed once with 150 uL PBS, and then lysed by a 5-minute incubation with with 50 uL iScript RT-qPCR Sample Preparation Reagent (BioRad) (64), containing the RNA spike-in control added at a final concentration of 2 pM,. We use 1 uL of this lysate in a 9 uL cDNA synthesis reaction, using the iScript cDNA Select Synthesis kit according to manufacturer’s instructions for synthesis with a gene-specific primer that binds immediately upstream of the barcode region (*cDNA_Fwd* 5’ *CTCCCTACAATGTCGGATTTGTATTTAATAG*-3’).

The resulting cDNA product was then amplified and the Read 2 sequence was added with PCR, using a forward primer that binds immediately upstream of the barcode, and appends the Read 2 sequence (*Rnd1_Fwd* 5’-*GTGACTGGAGTTCAGACGTGTGCTCTTCCGATCTCTCCCTACAATGTCGGATTTGTATTTAATAG* -3’) and a reverse primer that binds to the terminal sequence of the vRNA (*Rnd1_Rev* 5’-*AGTAGAAACAAGGGTGTTTTTCCTTATATTTCTGAAATCC*-3’), in a 50 uL reaction, with 5 uL of the cDNA reaction as template, according to manufacturer’s instructions, using KOD Polymerase Hot Start 2x Mastermix (Sigma). This amplification was performed with 20 cycles total, using an annealing temperature of 56°C, and elongation time of 20 seconds, ending with a final elongation step at 70°C for 2 min. A second round of PCR with unique dual indexing primers was performed to uniquely label each sample, using primers designed as follows UDI_i7 5’-*CAAGCAGAAGACGGCATACGAGATnnnnnnnnnnGTGACTGGAGTTCAGACGTGTGCTCTTCCGATCT-3* and *UDI_i5 5’-AATGATACGGCGACCACCGAGATCTACACnnnnnnnnnnACACTCTTTCCCTACACGACGCTCTTCCGATCT*-3’, using a 20-cycle protocol with KOD Polymerase Hot Start 2x Mastermix (Millipore Sigma), 20 seconds at 95°C, 10 seconds at 66°C and 20 seconds at 70°C, and a final 2-minute elongation at 70°C. After round 2 indexing PCR, samples were pooled at equal volume, run on a 1% agarose gel at 85V for 40 minutes. The band was extracted and purified with Nucleospin Gel Extraction Kit (Takara). The gel-purified DNA was then further purified with Ampure XP beads (Beckman Coulter) by adding 2x volume of DNA binding beads to the sample, incubating 5-10 minutes, then applying to a magnet and incubating again for 10 minutes, removing the supernatant, washing the pellet 2x with 80% ethanol, drying for approximately 10 minutes, taking care not to overdry the pellet, and then resuspending in elution buffer (5 mM Tris/HCl, pH 8.5).

The indexed DNA was quantified with the Qubit™ 1X dsDNA High Sensitivity Kit (Thermo Scientific), diluted to 4 nM and submitted for Illumina Sequencing on the NextSeq as either P1 or P2 depending on whether 1 or 2 plates were sequenced, to give an average number of reads per well of approximately 500,000-1,000,000 reads per well. This read depth is larger than necessary, but it allows for variability in sample loading in the absence of more rigorous, concentration-dependent pooling, which may introduce additional errors given the number of samples pooled per run. We use a 50 bp read length, and custom UDI indexes based on the sequences from Twist Biosciences. We identified an issue in sequencing of one of the spike-in control barcodes that began with “GG”.

Low read counts were detected for this barcode in a consistent well on each plate, though the affected well was different between plates that used either index set A or index set B, indicating that the index sequence may play a role in whether this occurs. The affected barcode was excluded from analysis. To eliminate this issue, barcodes that start with “GG” should be avoided in library design, and indices starting with “GG” should be also avoided (as recommended by Illumina).

### Sequencing-based neutralization assay plate setup and workflow

A step-by-step protocol is available on protocols.io (https://dx.doi.org/10.17504/protocols.io.kqdg3xdmpg25/v1). Briefly, the setup for a sequencing-based neutralization assay resembles other 96-well plate neutralization assays (Fig. 1). Prior to performing the assays, serum was treated with receptor destroying enzyme (RDE) and heat inactivated. RDE treatment was done to remove sialic acids in the serum which may interfere with the assay. This protocol was performed as previously described (35). To set up sequencing-based neutralization assays, virus and RDE-treated serum aliquots were thawed at room temperature. Then serial dilutions of serum were made in Influenza Growth Medium in a 96-well plate for a final total volume of 50 uL at 2x of the intended serum concentration to be incubated with virus. For this study, we performed 3-fold dilutions of serum with a starting concentration of 1:20 to capture a large range of potential NT50 values. Pooled virus library was diluted to an appropriate concentration, as determined by transcriptional titering, in Influenza Growth Medium (for this library, 70 uL of the stock library in 5430 uL of IGM) and 50 uL of diluted virus library was added to plate (Fig. 1). Virus-serum mixtures were incubated at 37°C with 5% CO2 for 1 hour. Following incubation of virus and serum, 50 uL of MDCK-SIAT1 cells at 1e6/mL in Influenza Growth Media were added to the plate, and the plate was then incubated at 37°C with 5% CO2 (Day 1, Fig. 1). At approximately 16 hours post infection (Day 2, Fig. 1), supernatant was removed, infected cells were washed once with 150 uL PBS to remove any residual virus. Cells were lysed with 50 uL of iScript RT-qPCR Sample Preparation Reagent (BioRad) per well containing the RNA spike-in control at 2 pM final concentration (Day 2, Fig. 1), cDNA for the barcoded region of the vRNA was synthesized and PCR was performed to prepare samples for Illumina sequencing as described in detail above.

### Participants and selection of samples

To illustrate the potential this new sequencing-based neutralization assay, we selected human serum samples from a randomized vaccine trial of repeated annual vaccination of Flublok done in Hong Kong in 2020-2025 (DRIVE-1 study) for testing. The DRIVE-1 study (ClinicalTrials.gov: NCT04576377) is a randomized placebo-controlled trial of repeated annual influenza vaccination in adults 18-45 years of age. We enrolled participants from the community if they met eligibility criteria and did not meet exclusion criteria. Individuals were eligible to participate if they were between 18 and 45 years of age, capable of providing informed consent, and intending to reside in Hong Kong for at least the next two years. Potential participants were excluded if they had received influenza vaccination in the preceding 24 months, if they were included in a priority group for influenza vaccination such as being a healthcare worker, if they had a diagnosed immunosuppressive condition or were taking immunosuppressive medication, or if they had severe allergies or bleeding conditions that contraindicated intramuscular influenza vaccination. We enrolled 447 participants between 23 October 2020 and 11 March 2021, and randomly allocated them to five separate groups, with annual receipt of vaccination (V) or saline placebo (P) as follows: V-V-V-V-V, P-V-V-V-V, P-P-V-V-V, P-P-P-V-V, or P-P-P-P-V. For the present study, we selected samples provided by 15 participants from year 1 and 2 of the V-V-(V-V-V) and P-V-(V-V-V) groups, i.e. participants randomly allocated to receive vaccination in both years, or placebo in year 1 (2020-2021) and vaccination in year 2 (2021-2022). The vaccine used was Flublok, a quadrivalent recombinant-HA vaccine produced in insect cells, including 45 mcg HA of each of four included strains as recommended by the World Health Organization for the northern hemisphere in 2020-2021 and 2021-2022 (65,66). The relevant H1N1 vaccine strains in 2020-2021 and 2021-2022 seasons were A/Hawaii/70/2019 (H1N1) (clade 5a.1) and A/Wisconsin/588/2019 (H1N1) (clade 5a.2). We collected 9ml sera immediately before administration of vaccine/placebo, and again at scheduled follow-up visits after approximately 30 days and 182 days each year. The study protocol was approved by the Institutional Review Boards of the University of Hong Kong (ref: UW19-551) and of the University of Chicago Biological Sciences Division (ref: IRB20-0217). Written informed consent was obtained from all participants. For the present analysis, we selected day 0, 30 and 182 sera collected from study year 2 in a subset of participants who had a pre-year 1 vaccination HAI titer <10 against the A/Hawaii/70/2019 (H1N1) vaccine strain , and who were assigned to either receive the placebo or the Flublok Vaccine (A/Hawaii/70/2019) in study year 1 (2020-2021 season) but all received the Flublok Vaccine (A/Wisconsin/588/2019) in year 2 (2021-22 season), i.e. two groups of participants who were vaccinated in 2021-2022 and who had either (1) prior year vaccination in 2020-2021 (“2xVax”) or (2) not (“1xVax”).

### Data analysis and computational pipeline

To analyze the data from the sequencing-based neutralization assay, we developed the modular *seqneut-pipeline* (https://github.com/jbloomlab/seqneut-pipeline),which we have made publicly available so that it can be easily used by others. This pipeline in turn makes use of the *neutcurve* Python package (https://github.com/jbloomlab/neutcurve) that was previously developed by some of us to automate the process of fitting neutralization curves. All analyses performed for this paper, which make use of *seqneut-*pipeline and *neutcurve* are available https://github.com/jbloomlab/flu_seqneut_DRIVE_2021-22_repeat_vax. We have also made available at https://jbloomlab.github.io/flu_seqneut_DRIVE_2021-22_repeat_vax/ interactive HTML versions of key plots and analysis notebooks to enable easier exploration of the data.

The initial processing of sequencing-based neutralization assay data is done by plate, as controls are included on each plate. First, we process Illumina barcode sequencing using the parser https://jbloomlab.github.io/dms_variants/dms_variants.illuminabarcodeparser.html to determine the counts of each barcoded strain in well of the 96-well plate in each selection condition. We then determine the average counts per barcode in each well, and filter wells on the plate with insufficient barcode counts. In addition, we examine the fraction of reads from the RNA spike-in barcodes and filter any wells with too few of these barcodes. Next, check that the fraction of counts from each barcode is consistent across all no-serum control wells and exclude any barcodes with highly variable representation in multiple control wells. See the *config.ym*l in the main project GitHub repo (https://github.com/jbloomlab/flu_seqneut_DRIVE_2021-22_repeat_vax) for the exact values used in these quality control steps.

We then calculate a fraction infectivity (F_b,s_) for each viral barcode (b) in each sample (s) as follows: F_b,s_ = [c_b,s_ / Σ_n_ c_n,s_] / [median_s0_(c_b,s0_ / Σ_n_ c_n,s0_)] where c_b,s_ is the count of barcode b in sample s, Σ_n_ c_n,s_ is the sum of the counts for all neutralization standard barcodes (n) in sample s, c_b,s0_ is the counts of viral barcode b in the no-serum sample (s0), and median_s0_(c_b,s0_ / Σ_n_ c_n,s0_) is the median taken across all no-serum control samples for the counts of the viral barcode divided by the total counts of all neutralization standard barcodes.

There is some variability in the amount of vRNA each influenza-infected cell produces (13,59). In general, this variability is averaged out by the fact that each barcoded virus infects hundreds of cells in our assay setup; nonetheless, rarely, a single barcode may be over-represented in a particular well of the plate. We can detect this issue by examining fraction infectivity for each barcode and excluding measurements where a given barcode in single well yields an extremely high fraction infectivity (such as a value > 5). After filtering is completed, we clip all fraction infectivity values at an upper limit of one, and then fit a Hill curve for each barcoded virus (Fig. 1), dropping any curves with a very poor coefficient of determination (r^2^). Again, see the *config.yml* for the exact filtering criteria used in this study.

Next, we calculate a neutralization titer for each barcoded strain. An NT50 can be calculated either by determining the midpoint of the fraction infectivity curve or directly calculating the value at which the fraction infectivity would be 50%. NT50s obtained by these two methods are generally very well correlated for each viral barcode; however, for curves with an upper plateau that is less than one (which sometimes happens if there is slightly lower representation of the RNA spike-in control barcode in the serum dilution wells than the no-serum control wells), then midpoint measurement may be a better representation of neutralizing titer. For this reason, we used the midpoint to calculate NT50s for this study.

Finally, we report the NT50 for each viral strain by as the median of the NT50s measured for each barcode corresponding to that strain. In instances in which there is a specific viral barcode with a NT50 that is an outlier with respect to the median, we manually examined the curves for these barcodes and excluded from the NT50 calculations if there were aberrant for an identifiable reason. At least two barcodes per strain must pass all filters for a NT50 to be reported in this study. See https://jbloomlab.github.io/flu_seqneut_DRIVE_2021-22_repeat_vax/ to visually inspect all neutralization curves generated for each plate and serum in this study.

### Traditional fluorescence-based neutralization assay used for validations

A traditional one-virus versus one-serum neutralization assays was used for the validations in Fig. 4. These assays were performed as previously described (21,22,67) (see https://github.com/jbloomlab/flu_PB1flank-GFP_neut_assay ) with the modifications described here. Briefly, viruses for neutralization assays were generated by reverse genetics as described above, using a single barcoded HA per strain. In place of the plasmid encoding the PB1 segment of A/WSN/1933, a plasmid (*pHH-PB1flank-eGFP)* which encodes GFP within the flanking sequences from the PB1 segment to allow for packaging into virions was used (21). These viruses were produced and passaged in cells constitutively expressing the PB1 protein (68). For reverse genetics, 5e5 293T-CMV-PB1 and 5e4 MDCK-SIAT1-CMV-PB1-TMPRSS2 cells were seeded in D10 media in a 6-well plate (67). Approximately 20 hours after seeding, cells were transfected with 250ng of each reverse genetics plasmids using BioT transfection reagent according to manufacturer’s instructions. At 20 hours post-transfection, D10 media was removed, cells were washed once with PBS and 2mL NAM (Neutralization Assay Media, consisting of Medium-199 supplemented with 0.01% heat-inactivated FBS, 0.3% BSA, 100 U/mL penicillin, 100 μg/mL streptomycin, 100 μg/mL calcium chloride, and 25mM HEPES) was added to the well. At approximately 48 hours post-transfection, the transfection supernatants were harvested, clarified by centrifugation at 845 x*g* for 4min, aliquoted, and frozen at -80°C. Viruses were passaged at low MOI (0.02) in MDCK-SIAT1-CMV-PB1-TMPRSS2 cells, as previously described. We titered the passaged virus stocks by TCID50 in 1e5 MDCK-SIAT1-CMV-PB1-TMPRSS2 (67). In addition, a test was performed to determine the amount of virus to add to each well such that the fluorescence signal decreases 2-fold with a 2-fold dilution of virus stock. Once the appropriate concentration of virus was determined, RDE-treated serum was serially diluted in a 96-well plate in NAM and virus was added. The viruses and serum were incubated at 37°C for 1 hour to allow antibodies in the serum to bind to GFP encoding influenza viruses containing the HA segment of interest and then 5e4 MDCK SIAT1-CMV-PB1 cells were added to each well. At approximately 20 hours post-infection, fluorescence was measured using a Tecan M1000 plate reader. Fraction infectivity was calculated as compared to a no-serum well. Neutralization curves were then fit to the data using the neutcurve package (https://jbloomlab.github.io/neutcurve).

### Hemagglutinin inhibition assay

The HAI titers reported in Fig. 4C were measured using protocol as described in (69). Briefly, serum samples were treated with receptor-destroying enzyme for 18 hours and heat-inactivated at 56°C for 30 min. Serum samples were serially diluted two-fold with a starting dilution of 1:10 and incubated for 30 minutes with four agglutinating doses (4AD) of test antigens. Titers were read as the reciprocal serum dilutions that inhibited agglutination of 0.5% turkey red blood cells.

---

**Figure S1.**
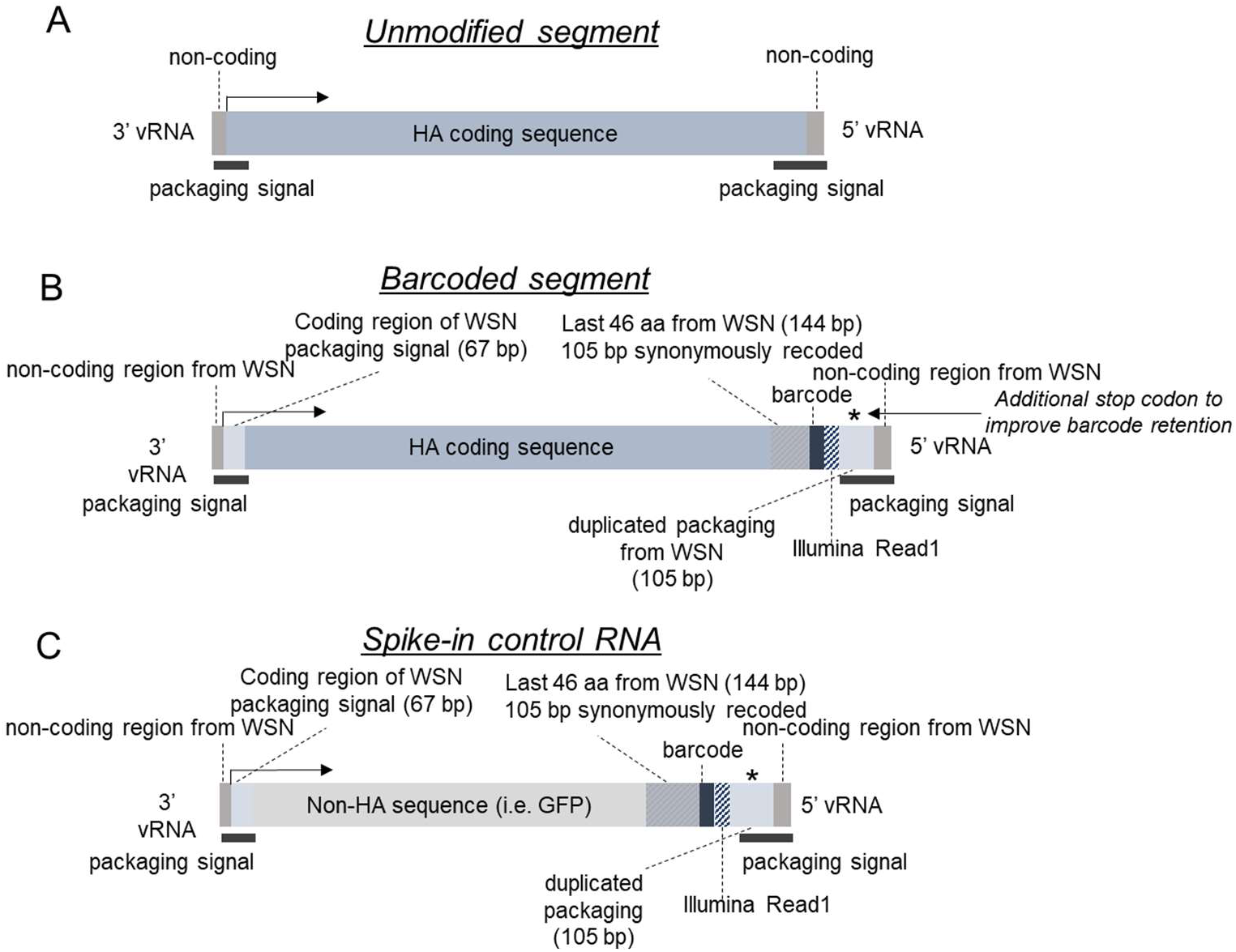
Design of barcoded HA genomic segment. A) An unmodified HA genomic segment with labels indicating the coding sequence, non-coding termini, and the packaging signals which span the non-coding regions into the coding sequence. Note that influenza viral RNA (vRNA) are negative sense, and the genomic segment is shown in the coding orientation so that the 3’ end of the vRNA is at left. We used a definition of the packaging signals spanning the first 67 nucleotides of the N-terminal region of the coding sequence (3’ end of vRNA) and the last 105 nucleotides of the C-terminal region of the coding sequence (5’ end of vRNA). B) The barcoded HA genomic segments are created by making several modifications. First, the noncoding region at the 3’ end of the vRNA as well as the first 67 nucleotides of the coding sequence are replaced with those of the lab-adapted A/WSN/1933 (H1N1) strain. Therefore, the full signal peptide from WSN HA is used in the final constructs. Second, the transmembrane domain and cytoplasmic tail are replaced with the homologous 46 amino-acid region from the WSN strain followed by a pair of consecutive stop codons (144 nucleotides, stop codons not indicated in schematic), with synonymous mutations introduced to remove the packaging signals without altering the encoded protein in the last 105 nucleotides of this region. This synonymous re-coding is to reduce homology between the duplicated packaging sequence within the coding region and the terminal packaging sequence used for incorporating the segment into the virion, which may be important for reducing intra-segment recombination. A 16-nucleotide barcode is placed immediately after the double stop-codon at the end of the coding sequence followed by an Illumina read 1 sequence. Finally, downstream of the coding sequence is the duplicated packaging signal region of the WSN virus (spanning the last 105 nucleotides of the coding sequence) followed by the 5’ noncoding region of the WSN viral RNA (45 nucleotides). A stop codon is engineered into the coding region of the duplicated WSN packaging signal so if there was homologous recombination between the duplicated region that removed the barcode, the resulting HA protein would be truncated and so non-functional. Plasmid maps for the barcoded library strains are available at https://github.com/jbloomlab/flu_seqneut_DRIVE_2021-22_repeat_vax/tree/main/plasmids C) The spike-in control RNA is generated to resemble the barcoded HA segment, with the 3’ packaging region from WSN, followed by a GFP sequence, the synonymously recoded transmembrane and c-terminal domain from WSN, a 16-nt barcode, an Illumina read 1 sequence, and the 5’ packaging sequence from WSN.

**Figure S2.**
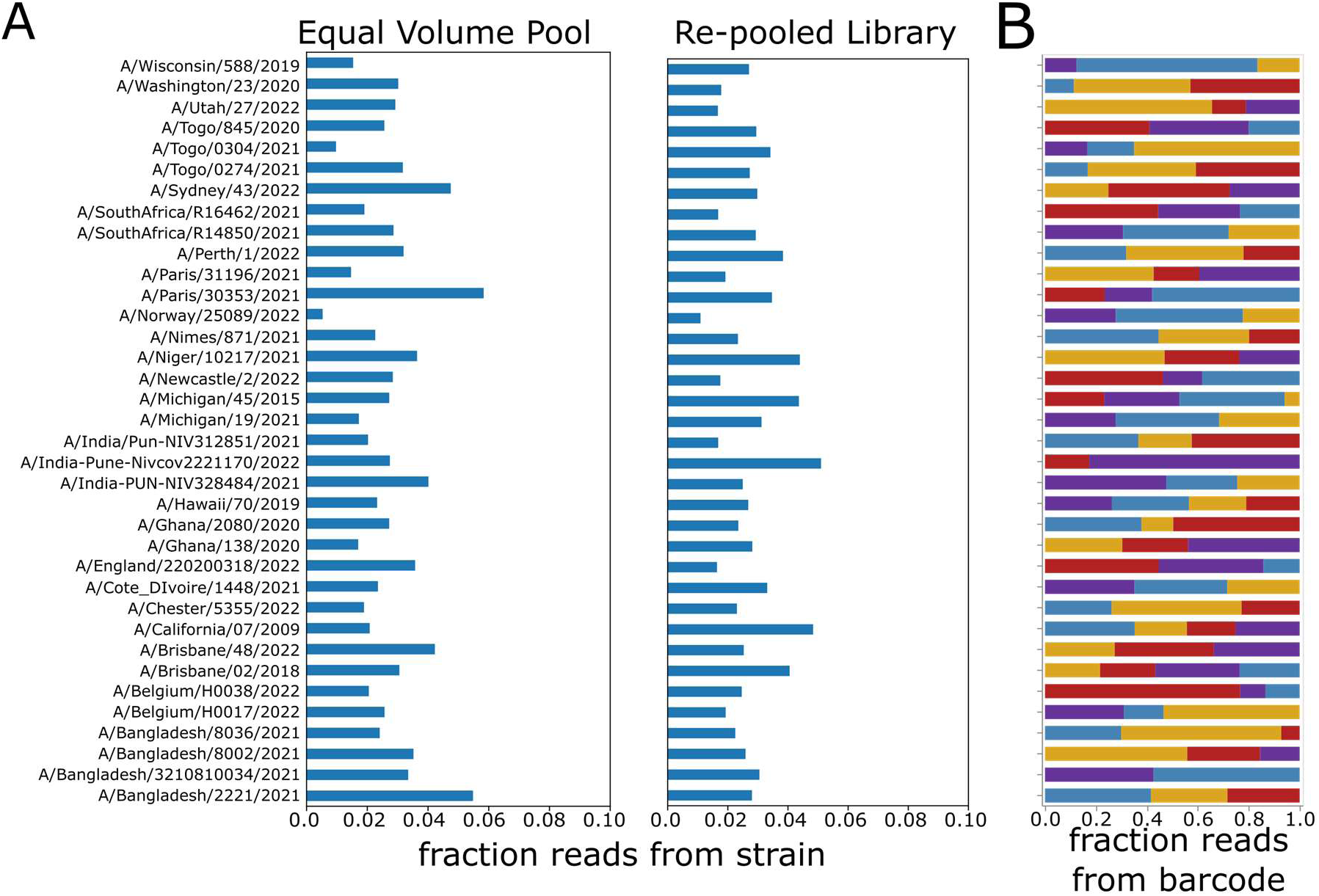
Sequencing-based pooling of library to try to achieve equal transcriptional titers across viruses. A) We first infected cells with equal volumes of the stocks of the viruses with each of the 36 different HAs in the library, and then sequenced the viral barcodes to quantify the fraction of barcodes attributable to each virus, as plotted at left. We then re-pooled the virus stocks to try to achieve equal barcode representation for each virus stock. The plot at right shows the fraction of barcodes attributed to each virus after the re-pooling. This re-pooled mix was used in all experiments in this study. C) The fraction of barcode reads attributable to each of the two to four unique barcodes for each virus in the re-pooled libraries. Each row represents a different virus strain and each color represents a different barcode, with the width of each stacked bar representing the fraction of counts for that strain that are attributable to each of the two to four barcodes for the strain.

**Figure S3.**
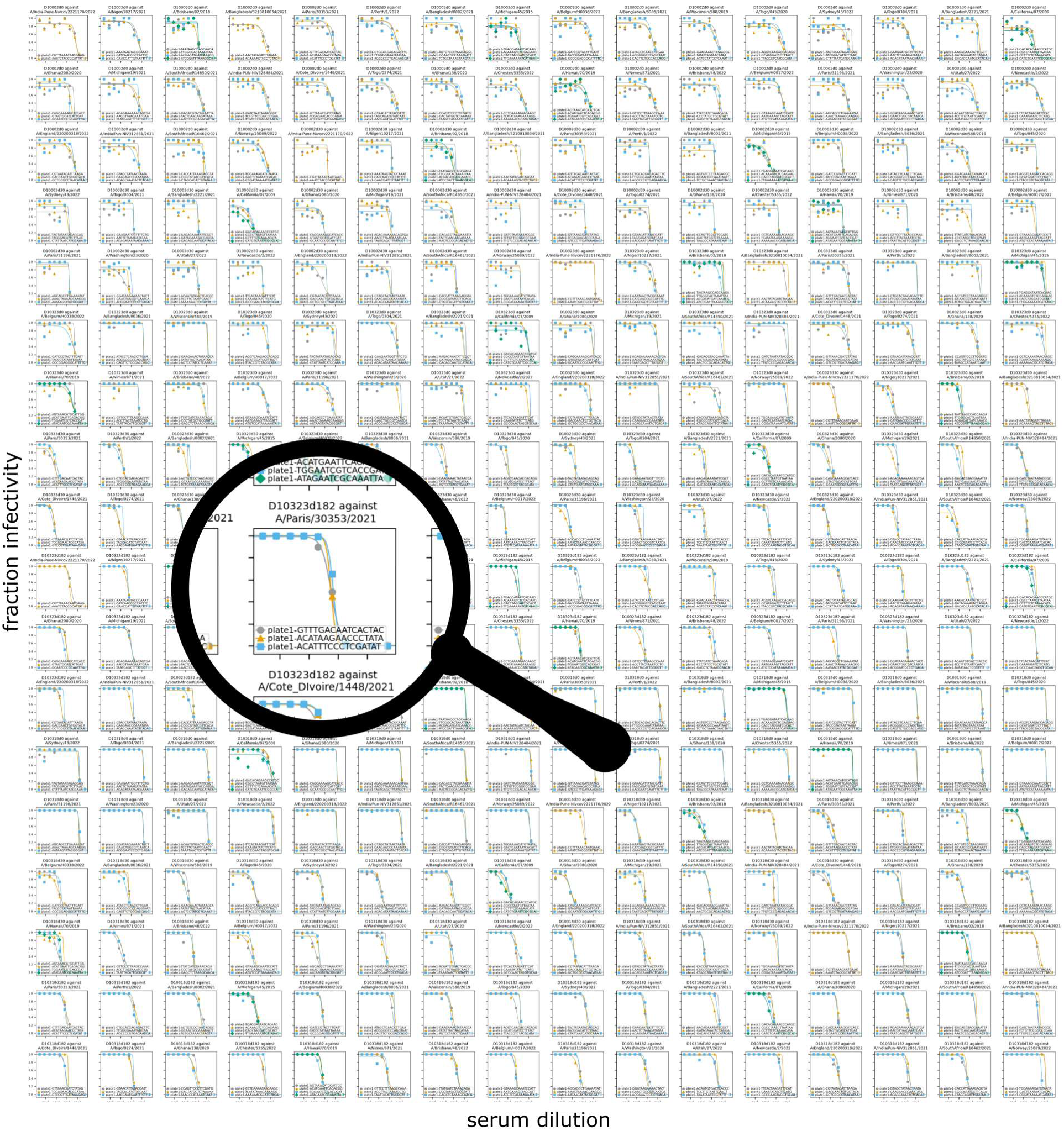
All neutralization curves generated on a single sequencing-based neutralization assay plate. Each plate yields neutralization curves for each of the two to four barcoded variants of each of the 36 HAs in the library against each serum sample run on the plate., This figure shows 288 plots each corresponding to 36 viral strains against 8 serum samples (each serum was diluted across one of the plate’s eight rows). A magnifying glass icon is used to show a single plot from this plate, which shows all three barcodes for the A/Paris/30353/2021 HA against serum D10323d182. Go to https://jbloomlab.github.io/flu_seqneut_DRIVE_2021-22_repeat_vax to see the full neutralization curves for all plates run in this study; for instance, see https://jbloomlab.github.io/flu_seqneut_DRIVE_2021-22_repeat_vax/process_plate1.html for the curves shown here.

**Figure S4.**
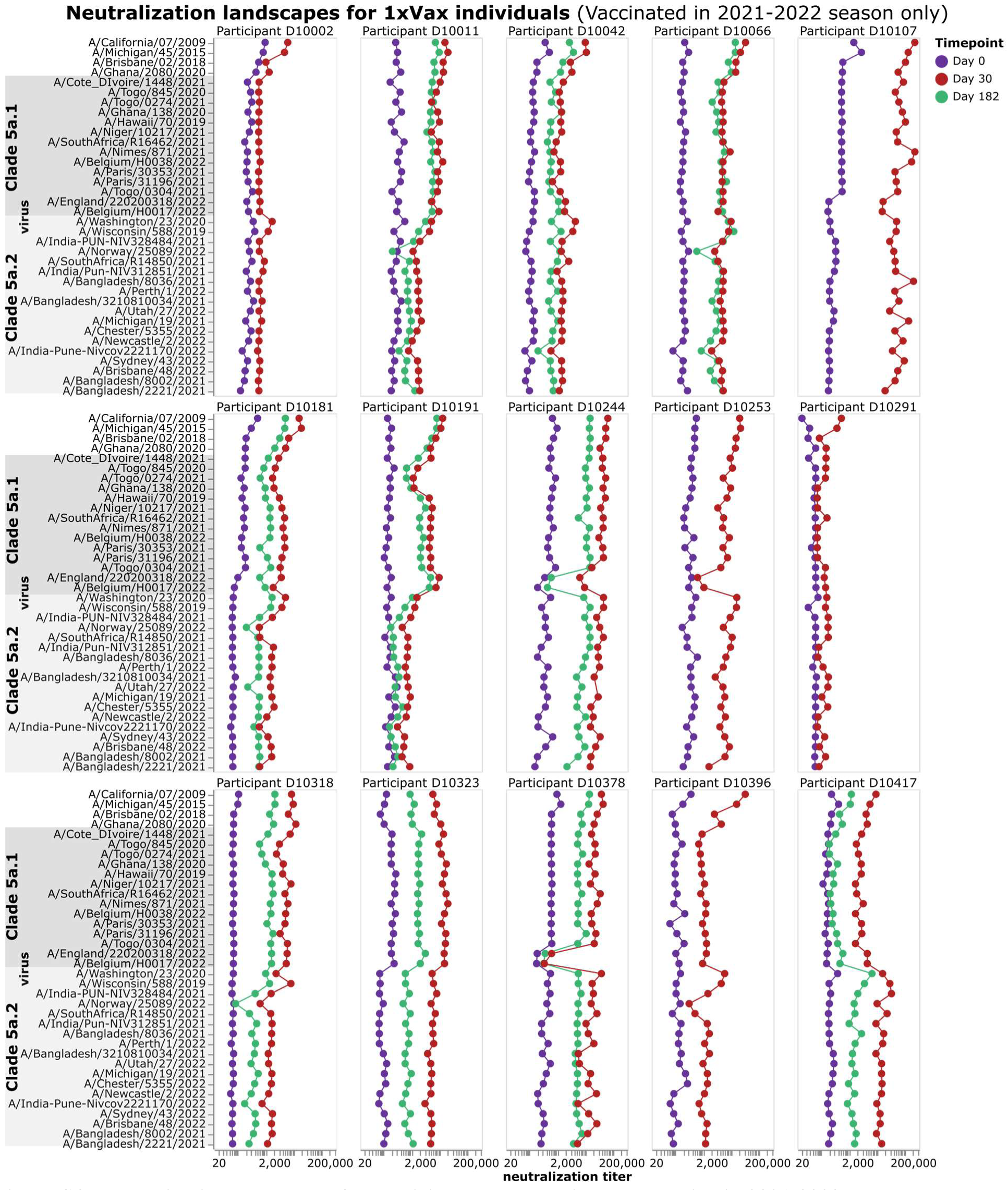
Neutralization landscapes for participants that received the vaccine in 2021-2022 season, but were not vaccinated the previous year.

**Figure S5.**
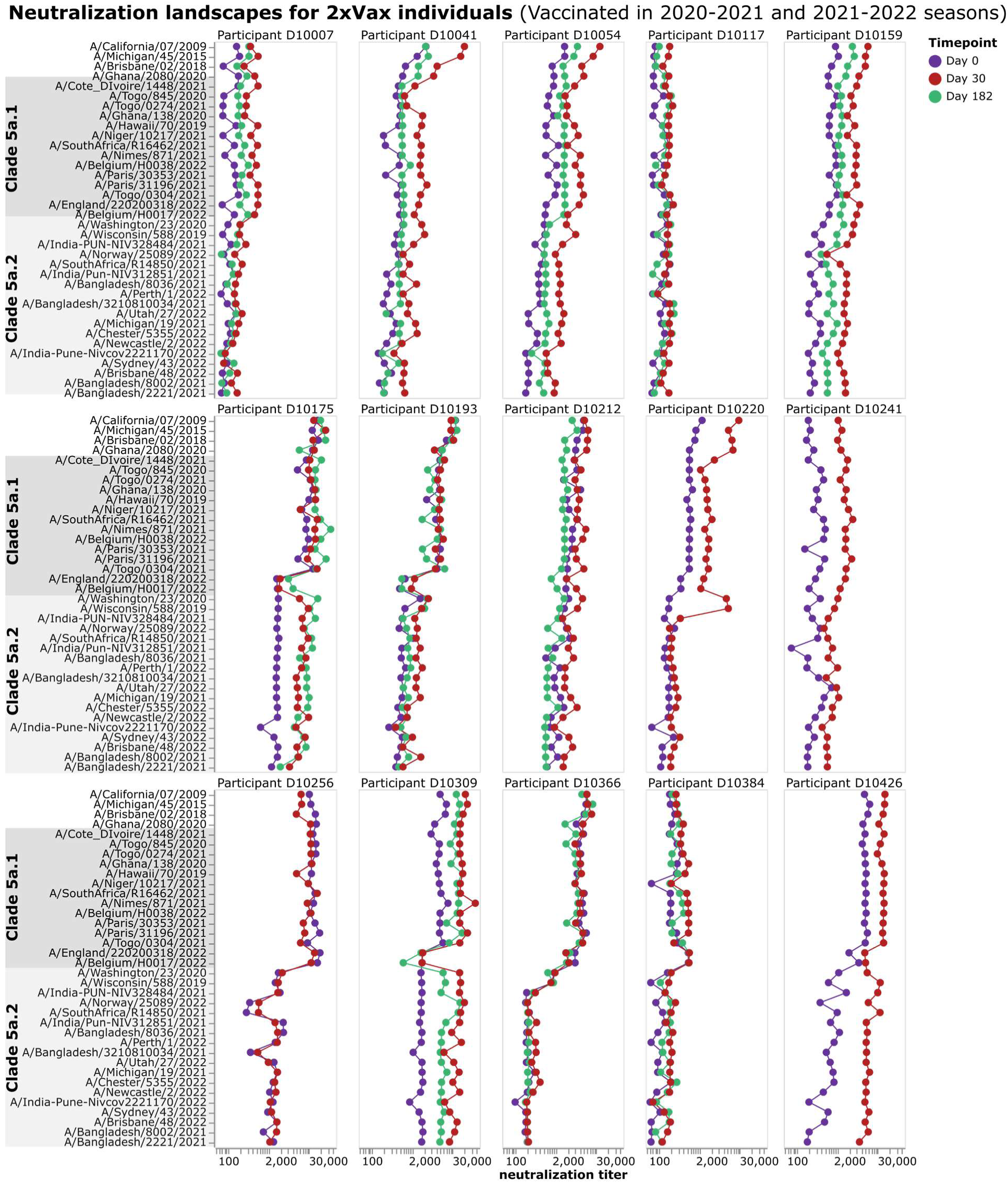
Neutralization landscapes for participants who received the vaccination in 2021-2022 and had also been vaccinated in the previous season.

**Figure S6.**
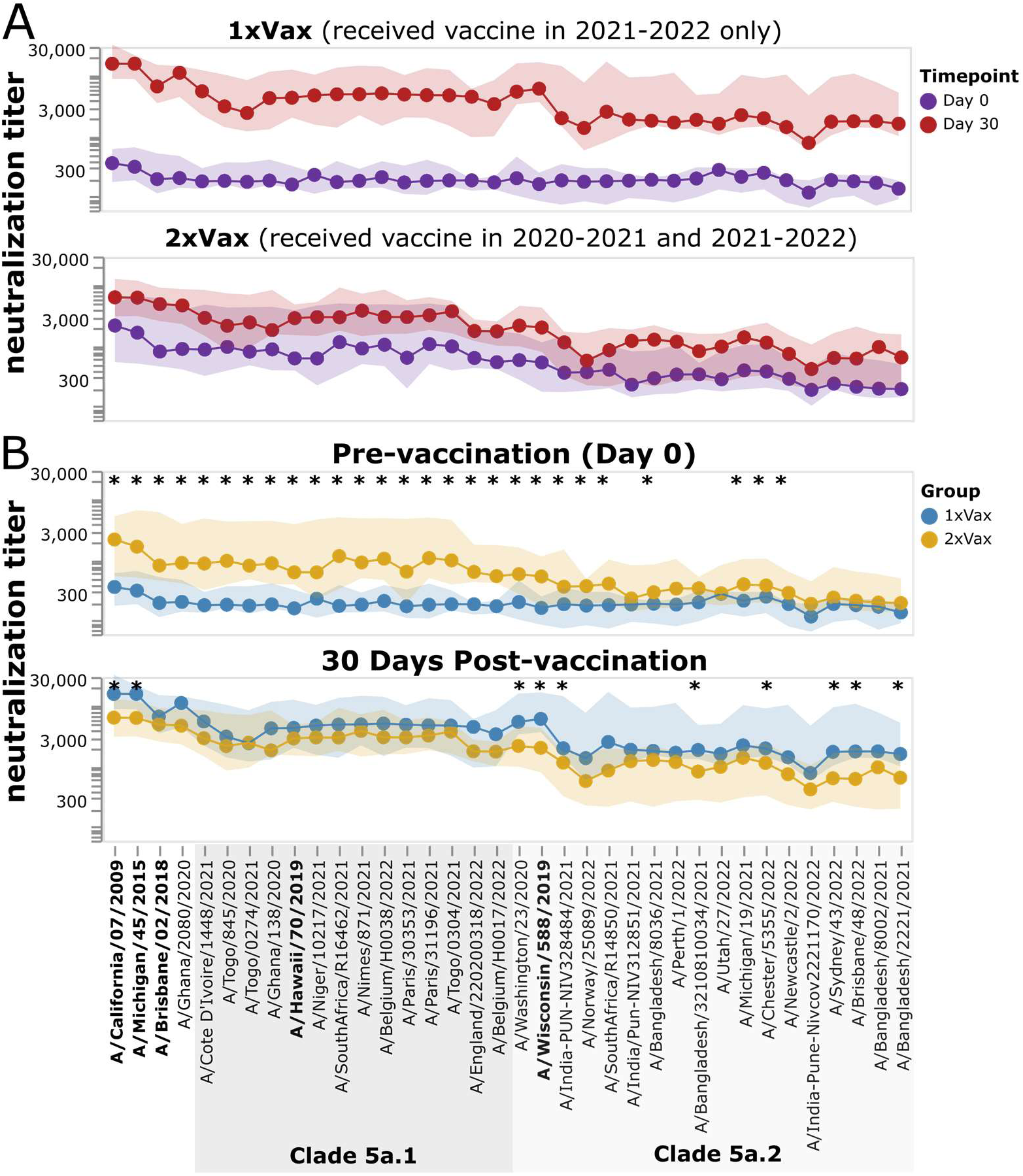
Neutralization titers of all study participants including those without a day 182 sample. This figure is comparable to Figure 6A and Figure 7 except it shows all study participants, whereas those other two figures only show participants who had a day 182 as well as a day 0 and day 30 sample. See Table 1 for the number of participants in each category. Strains with a significant difference in median titer between groups as assessed by a Mann-Whitney U Test are indicated with an asterisk at the top of each plot panel.

**Figure S7.**
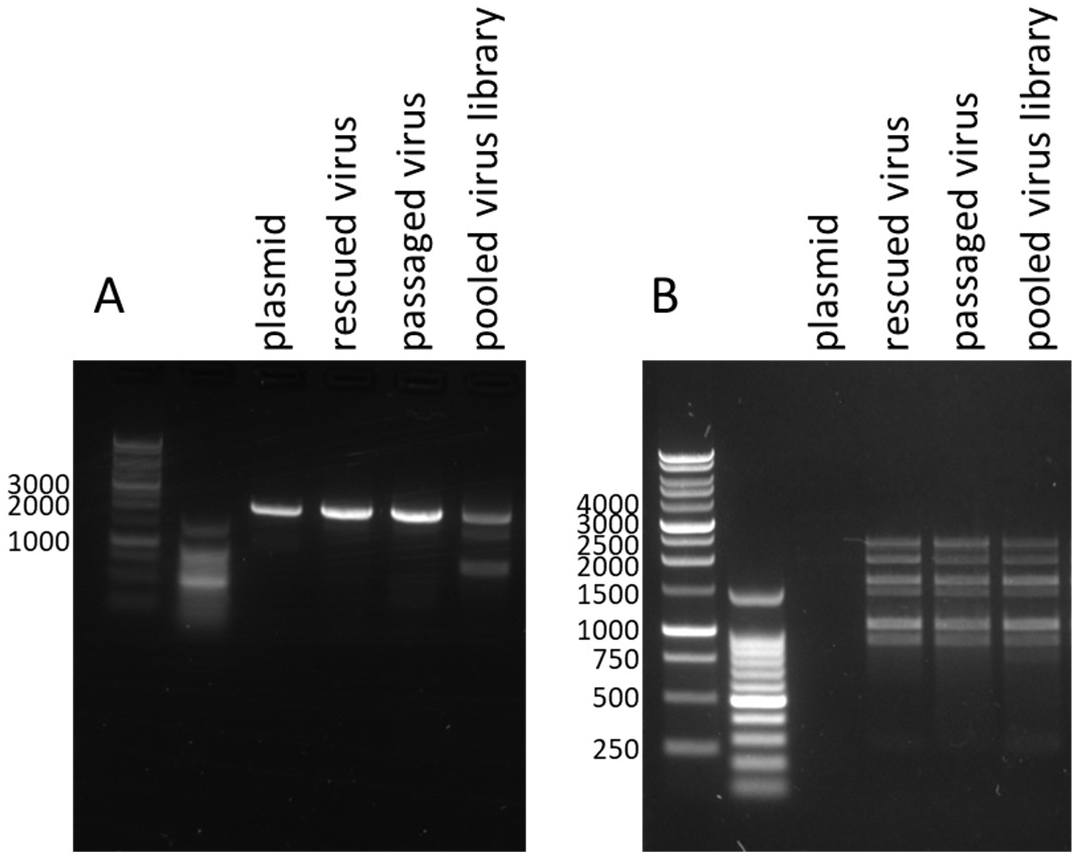
The HA barcode region is stably retained, and the viral stock is relatively free of high-abundance internal deletions in the viral gene segments. A) Full-length cDNA synthesis and PCR amplification of the HA genomic segment for a single barcoded plasmid control, a reverse-genetics rescued barcoded strain, a passaged barcoded strain, and the pooled barcoded library. Only the full-length HA band is observed for the rescued and passaged single strain, although some shorter deletion products are found in the pooled library. These bands were verified by Sanger sequencing to contain the barcode region (additional details regarding these products in the methods section). B) Full-length cDNA synthesis and PCR of all viral genomic segments for a single barcoded plasmid control, a reverse-genetics rescued barcoded strain, a passaged barcoded strain, and the pooled barcoded library. No band is present in the plasmid control, likely as the PCR conditions needed to amplify off of a circular segment differ from those needed to amplify from purified reverse-transcribed linear viral cDNA. Bands are present for all expected sizes of influenza genomic segments in RNA extracted from the viral supernatant, with no especially high intensity shorter product bands observed, indicating that the viral stock is not overly dominated by defective particles with internal deletions in the genes.

